# A Chromatin-Structure-Guided Framework for Predictive and Interpretable Regulatory Genomics

**DOI:** 10.1101/2025.11.03.686435

**Authors:** Bowei Ye, Lin Du, Min Chen, Yang Dai, Ao Ma, Jie Liang

**Affiliations:** Center for Bioinformatics and Quantitative Biology, and Richard and Loan Hill Department of Bioengineering, University of Illinois Chicago, Chicago, IL 60607, USA; State Key Laboratory of Systems Medicine for Cancer and Bio-ID Center, School of Biomedical Engineering, Shanghai Jiao Tong University, Shanghai 200240, China; University of Illinois Cancer Center, Chicago, IL 60612, USA

## Abstract

Chromatin organization shapes gene regulation by linking distal elements across megabase scales, yet most predictive genomics models still treat the genome as linear, without incorporating three-dimensional structure. Hi-C provides genome-wide chromatin conformation information, but its contact maps are population-averaged, distance-biased, and noisy, obscuring the biologically specific contacts. We present CHROME, a framework built on a self-avoiding polymer ensemble null model that identifies physically specific, non-random Hi-C contacts. By integrating these contacts into graph representations, CHROME enables efficient information transfer across spatially connected loci. It integrates sequence, chromatin accessibility, or pre-trained embeddings into a graph attention architecture to predict cell line-specific ChIP-seq profiles, consistently outperforming local encoder baselines and generalizing to an unseen cell line. The resulting graph embeddings also enhance prediction on tissue-specific eQTL and ClinVar variant pathogenicity, outperforming local sequence-based embeddings. Beyond predictive performance, CHROME provides interpretability through attention-derived neighbor-to-center contributions that reveal how spatially connected loci influence local regulatory activity over multi-megabase distances. Together, these results show that incorporating physically validated chromatin interactions enables more accurate and interpretable modeling of gene regulation and variant effects.

## 1 Introduction

The regulation of gene expression and DNA replication arises from a complex interplay among DNA sequence, chromatin state, and three-dimensional genome architecture (1–3). Advances in chromosome conformation capture technologies (4–6), particularly Hi-C [7, 8] data from the 4D Nucleome Project [9], have generated extensive maps of three-dimensional genome organization across diverse tissues and cell types.

Over the past decade, deep learning has transformed our ability to predict and interpret epigenomic signals, with models increasingly learning predictive features directly from raw genomic inputs. Early efforts such as DeepSEA (10–12) demonstrated that the convolution neural network [13] could predict a wide range of epigenetic profiles from DNA sequence alone. This was extended by variant-effect predictors like ExPecto [14], which estimated the functional impact of mutations. To capture distal regulatory interactions, models including Basenji [15] and the transformer-based Enformer [16] expanded receptive fields to enable long-range sequence modeling. Other sequence-based approaches, such as Sei [17], emphasized interpretability by grouping predictions into regulatory programs. Moving beyond sequence-only inputs, multi-modal frameworks such as EPCOT [18] combined sequence with chromatin accessibility to improve predictive power. More recently, large-scale models have pushed generalization further: EVO2 [19] generates universal genomic embeddings, Borzoi [20] and GET [21] predict RNA-seq tracks, and AlphaGenome [22] integrates epigenetic, transcriptomic, and Hi-C data, extending the receptive field to 1 Mb. Yet, scaling purely linear sequence-based architectures to multi-megabase contexts remains computationally demanding, limiting their ability to capture very long-range dependencies in practice. This motivates interest in alternative modeling paradigms that can efficiently represent extended genomic contexts.

Despite these advances, most existing approaches still treat the genome as a linear sequence, overlooking the three-dimensional chromatin organization revealed by Hi-C. Structural features such as topologically associating domains (TADs), compartments, and loops, often extending beyond 1 Mb, connect distal enhancers to promoters and align with cell-type-specific epigenetic profiles [23, 24], including ChIP-seq signals. Although Hi-C data capture these structural relationships, extracting biologically meaningful and physically specific, non-random contacts remains challenging because interaction frequencies are population-averaged, distance-biased, and affected by experimental noise [25, 26]. In contrast to the protein field, where topology has been extensively studied [27, 28] and graph-based models have proven effective (29–31), analogous graph frameworks for chromatin remain less developed. Existing methods such as GraphReg [32] incorporate Hi-C contacts into graph structures but restrict edges to interactions within 2 Mb, limiting their ability to capture broader long-range dependencies.

These limitations motivated the development of a deep learning-driven, chromatin-structure-guided framework that leverages the spatial information from Hi-C to integrate biologically meaningful contacts across multi-megabase scales for diverse predictive tasks. To extract such contacts from Hi-C data, we employed CHROMATIX [33], which identifies physically specific, non-random contacts using a self-avoiding polymer ensemble null model.

Building on these validated non-random contacts, we developed CHROME (CHROMatin-structure–guided graph Embedding framework), which incorporates non-random contacts into a unified graph representation that enables efficient information flow across distal regulatory elements. Although broadly applicable to various genomic prediction tasks, we demonstrate CHROME using ChIP-seq signal prediction as a representative case. CHROME introduces four key innovations: (i) Hi-C contact filtration using CHROMATIX [33], retaining only physically specific, non-random interactions; (ii) an extended receptive field of up to 4 Mb to capture long-range regulatory influences; (iii) flexible locus-level embeddings accommodating raw sequence, chromatin accessibility, or pre-trained representations; and (iv) intrinsic interpretability, with graph attention layers that assign neighbor-to-center contribution weights.

Applied to GM12878, IMR-90, and K562, CHROME consistently improved predictive performance for raw sequence, DNase + sequence, and Evo2 embeddings, yielding systematic gains over local encoder-only baselines. In all cases, locus-level representations from baseline encoders were contextualized by graph attention over non-random contacts. On an unseen cell line (HepG2) not included during training, CHROME maintained robust improvements, demonstrating strong cross-cell-type generalization. Beyond regulatory prediction, graph embeddings from trained CHROME models improved tissue-specific expression quantitative trait locus (eQTL) classification (mapping GM12878→lymphoblastoid [blood] and IMR-90→fibroblast [lung] tissues from GTEx [34]) and ClinVar variant pathogenicity prediction [35]. Because CHROME is trained on regulatory profiles, its attention-derived contribution maps provide direct interpretability, revealing how spatially connected, nonrandom loci influence local regulatory activity. These contributions remain detectable across multi-megabase distances, with ablation tests confirming a clear preference for non-random over random Hi-C contacts. Together, our findings show that physically validated chromatin contacts are functionally informative for gene regulation, positioning CHROME as a generalizable and interpretable framework that leverages three-dimensional genome structure to improve regulatory prediction and variant interpretation.

## 2 MATERIALS AND METHODS

CHROME is a chromatin-structure-guided graph embedding framework that integrates DNA sequence, chromatin accessibility, or pre-trained embeddings with CHROMATIX-validated [33] non-random chromatin contacts derived from a self-avoiding polymer ensemble null model (Figure 1), enhancing the prediction of target ChIP-seq profiles and providing interpretability by revealing how spatially connected, non-random contacts shape regulatory activity. In this framework, each signal peak is represented as a signal-centered subgraph, with the signal-associated locus as the center node and directly interacting non-random loci as neighbors. Locus-level features are encoded and propagated through graph attention networks, capturing information flow across both cis and distal, non-randomly connected loci. The detailed architecture and downstream tasks are described in the following sections, while the CHROMATIX procedure for identifying non-random contacts, along with dataset descriptions, hyperparameters, and training procedures, are provided in the Supplementary Information (Sections S1-S4). All study mapped to the hg38 reference genome.

**Figure 1:**
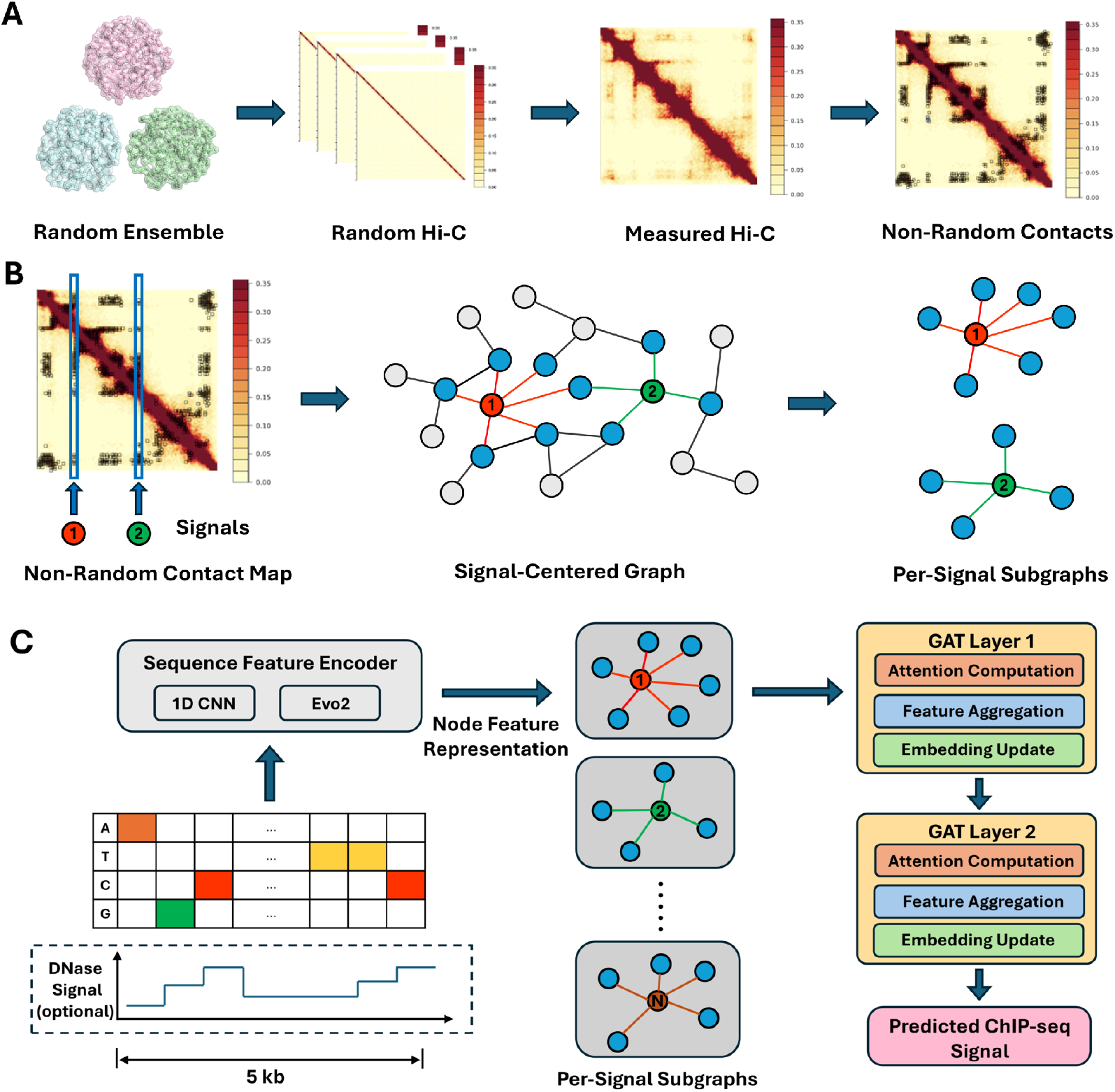
Chromatin-structure-guided graph framework (CHROME). (**A**) CHROMATIX validation process for Hi-C data. A self-avoiding polymer ensemble model generates random three-dimensional chromatin conformations within a confined nuclear volume, forming a physical null distribution of contact probabilities. Measured Hi-C contact frequencies are statistically compared against this null to identify physically specific, non-random contacts, which are subsequently used for CHROME graph construction. (**B**) Construction of signal-centered graphs. Non-random contact maps are overlaid with 5 kb loci containing ChIP-seq signals. Example signal loci are highlighted in red and green, serving as graph centers, with their direct non-random neighbors shown in blue and unconnected loci in gray. Each signal-centered subgraph captures the spatially connected regulatory context of its center locus, and overlaps may occur when nearby signals share neighbors. (**C**) CHROME architecture. Each 5 kb locus is encoded using a sequence feature encoder (1D CNN for one-hot DNA sequence or DNA sequence with DNase values, and MLP for Evo2 embeddings). Encoded nodes are assembled into signal-centered subgraphs, which are propagated through two graph attention (GAT) layers for attention-based feature aggregation and embedding update. The resulting graph embeddings are used to predict 751 ChIP-seq profiles in a cell line-specific manner.

### Graph from CHROMATIX-Validated Hi-C Contacts

We used bulk Hi-C data for GM12878 (4DNES3JX38V5), IMR-90 (4DNES1ZEJNRU), K562 (4DNESWST3UBH), and HepG2 (4DNFICSTCJQZ; chromosome 9 only), all obtained from the 4D Nucleome Consortium [9, 36]. To identify non-random contacts, for each cell line, 5 *×* 10^5^ self-avoiding polymer chains were simulated under confined nuclear volume following the CHROMATIX framework [33], which estimates null contact probability distributions from a physical polymer-ensemble model. Interactions passing a false discovery rate (FDR) threshold of 0.05 were retained as physically specific, non-random contacts, representing interactions unlikely to arise from random polymer folding and thus biologically meaningful. At 5 kb resolution, modeling chromatin chains of 4 Mb (800 loci) has been shown to capture the majority of TADs and enhancer-promoter hubs while remaining computationally feasible [37], thereby providing a practical upper bound for the receptive field considered in CHROME (Figure 1A). Across the whole genome, the proportion of non-random contacts identified was approximately 1.8% in GM12878, 6.0% in K562, and 5.0% in IMR-90 (representative 4 Mb regions and more detailed non-random contact analysis provided in Figures S1).

From the processed non-random contact maps, we constructed signal-centered subgraphs *G*_*s*_ = (*V*_*s*_, *E*_*s*_) for each ChIP-seq signal locus *s*. The node set *V*_*s*_ comprises genomic loci partitioned into non-overlapping 5 kb loci, and the edge set *E*_*s*_ includes direct, non-random contacts linking each signal center locus to loci with physically specific interactions. For each ChIP-seq peak defined at 1 kb resolution, a 5 kb window spanning 2 kb upstream and downstream flanking regions was defined as the center node. Each peak was thus represented as a signal-centered subgraph, with overlapping subgraphs sharing nodes when multiple peaks occurred in close proximity (Figure 1B).

### CHROME Framework for ChIP–seq Prediction

To model how non-random chromatin contacts influence regulatory activity, CHROME employs a graph attention framework [38] to predict cell line-specific ChIP-seq signals. Each signal-centered locus is represented as a node with sequence, chromatin accessibility, or embedding-derived features, while non-random Hi-C contacts define the edges through which information is propagated. This design enables the model to integrate information from both cis and distal, spatially connected non-random loci, thereby capturing cell line-specific transcription factor and histone modification patterns within a chromatin-structure-aware architecture.

*Node embeddings*. CHROME was trained on GM12878, IMR-90, and K562, predicting cell line-specific ChIP-seq profiles covering a total of 751 assays from ENCODE [39] (539, 168, and 44 profiles, respectively) derived from signal-centered subgraphs (Figure 1C). Model generalization was further evaluated on an unseen cell line, HepG2 (chromosome 9 only), which includes 324 additional profiles. Detailed assay information is provided in Figure S4. Each subgraph *𝒢*_*s*_ = (*𝒱*_*s*_, *ℰ*_*s*_) consists of a center node *c*, corresponding to the 5 kb bin containing the labeled 1 kb region, and its directly contacting non-random Hi-C neighbors *𝒩* (*c*) at 5 kb resolution. Every 5 kb locus *v* is represented by a *d*-dimensional embedding

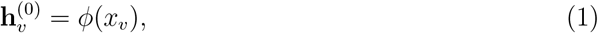

where *x*_*v*_ denotes the raw locus features. Because data availability and information content vary across modalities, we implemented three encoder variants to balance general applicability and predictive power: (i) The raw DNA sequence was one-hot encoded and processed by a 1D CNN [13] following the EPCOT pretraining design [18]. (ii) To incorporate chromatin accessibility, the base-resolution DNA one-hot encoding was concatenated with cell line-specific DNase signals and processed by the same CNN backbone. (iii) To evaluate the representational power of pretrained sequence models, the raw DNA sequence was embedded by Evo2 (7B) [19], producing a dense per-locus vector that was processed by a multilayer perceptron (MLP) [40]. All encoder variants map their inputs into a common *d*-dimensional space, which serves as the input for graph propagation.

#### Graph attention propagation

Information flows over non-random contacts using two layers (*l* = 0, 1) of graph attention [38]. At each layer, node features are first linearly projected, 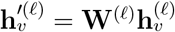. For a single head, attention logits between *v* and a neighbor *u* ∈ 𝒩 (*v*) are

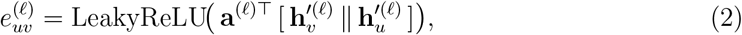

normalized by a softmax

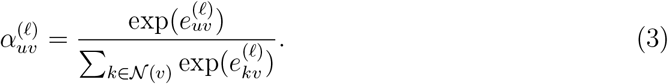

Node updates aggregate neighbor messages,

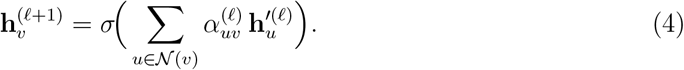

With *M* attention heads, the multi-head update is

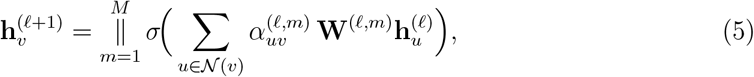

where ∥ denotes concatenation across heads. Residual connections, dropout, and normalization are applied after each layer. The coefficients 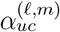 act as intrinsic attributions of neighbor *u*’s contribution to the center node *c* at head *m*, with message 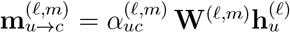.

#### Two-stage training

Training proceeds in two stages. In the first stage, we pretrain the locus encoders on center nodes only until validation performance saturates. For DNA sequence or sequence+DNase inputs, a 1D CNN processes 4*×*5000 or 5*×*5000 matrices, respectively. For Evo2 embeddings, a frozen 7B model produces 1*×*4096 vectors that are projected by a lightweight MLP [40]. Pretraining follows a chromosome split with chr9 held out for testing, chr8 for validation, and all remaining for training (chrY excluded).

In the second stage, the pretrained CNN or MLP weights initialize the encoders for all graph nodes. For each signal-centered subgraph, the center node’s input is passed through the encoder to obtain a center embedding 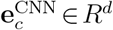. In parallel, all neighbor nodes are encoded by the same CNN/MLP (shared weights) to generate node features for GAT propagation. After two GAT layers[38], node embeddings are pooled with global mean over the subgraph to yield 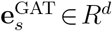. The final embedding is the concatenation

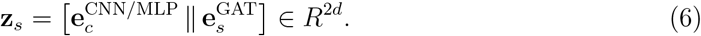

Fine-tuning is performed end-to-end under the same chromosome split, with early encoder layers frozen and later layers unfrozen. This regimen preserves low-level motif features learned during pretraining while allowing higher layers to adapt to graph-contextualized supervision.

#### Prediction objective and optimization

The concatenated embedding **z**_*s*_ *∈ R*^2*d*^ is passed through a linear classifier to produce logits

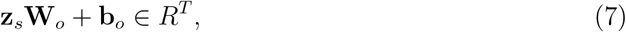

which are converted to probabilities

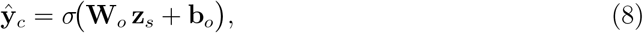

where *σ* is the elementwise logistic function and *T* =751 corresponds to the number of ChIP-seq profiles. Since each locus can be associated with multiple assays, training is formulated as a multi-label prediction problem. With ground-truth labels *y*_*c,t*_ *∈ {*0, 1*}* and predicted probabilities *ŷ*_*c,t*_, the per-center masked binary cross-entropy loss is

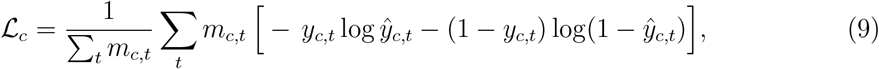

and the dataset objective is 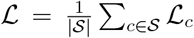. Here *m*_*c,t*_ ∈ {0, 1} is an availability mask ensuring that only observed assays contribute.

We implement this objective using BCEWithLogitsLoss, optimized with Adam (learning rate 3*×*10^*−*5^, weight decay 10^*−*6^) and a ReduceLROnPlateau scheduler (mode=min, factor=0.5, patience=1, min lr=10^*−*7^).

### Variant Analysis with CHROME

*eQTL prediction*. To evaluate how CHROME’s learned regulatory representations generalize to eQTL interpretation, we aligned cell lines to their most closely related GTEx [34] tissues: GM12878 was associated with Blood (*EBV-transformed lymphocytes*), and IMR-90 with Lung (*Cultured fibroblasts*). We adopted the same eQTL datasets as used in Borzoi [20], restricting analysis to these two tissues. For each candidate eQTL, we defined two inputs: (i) the 5 kb locus centered on the transcription start site (TSS), and (ii) the 5 kb locus containing the variant, together with its tissue (cell line)-specific non-random contacts.

The TSS-centered locus was processed by a pretrained locus encoder (CNN for DNA or DNA+DNase inputs; MLP projection for Evo2 embeddings), yielding a TSS embedding **e**_TSS_ *∈ R*^*d*^. In parallel, the variant-centered subgraph *𝒢*_*v*_ was propagated through the CHROME GAT trunk, producing a variant embedding **e**_var_ ∈ *R*^*d*^. The two embeddings were concatenated,

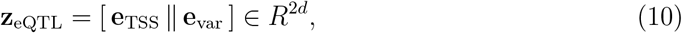

and passed to an MLP for binary classification of whether the variant corresponds to an eQTL. Training was performed using cross-entropy loss.

#### ClinVar variant pathogenicity classification

We further evaluated how the regulatory knowledge learned by CHROME generalizes to ClinVar variant pathogenicity [35]. Variants annotated as benign or pathogenic were collected, including both coding and noncoding entries. Because ClinVar annotations are not tissue-or cell line-specific, we constructed an averaged input representation: non-random contacts were intersected across GM12878, IMR-90, and K562 to define a consensus non-random contact map, and DNase accessibility was averaged across the same three cell lines to approximate a general open chromatin state. For each variant, the 5 kb locus containing the variant and its neighbors in the consensus non-random contact map were extracted to form a variant subgraph. Locus-level features (DNA sequence, DNA+DNase, or Evo2 embeddings) were encoded and propagated through GAT layers, producing a variant embedding **e**_ClinVar_ *∈ R*^*d*^. This embedding was then passed to an MLP for binary classification of benign versus pathogenic, trained with cross-entropy loss.

## 3 RESULTS

### Accurate Prediction of ChiP-Seq Profiles

We first evaluated CHROME on the prediction of cell line-specific transcription factor and histone modification profiles, a widely used benchmark in regulatory genomics for assessing whether non-random, spatially connected loci contribute additional information that improves predictive performance. This task requires integrating both sequence information and cell line-specific context, making it well-suited for a framework that incorporates cell line-specific non-random three-dimensional neighborhoods.

Across 751 profiles in GM12878, K562, and IMR-90, CHROME consistently outperformed local encoder-only baselines that used the same input features without neighborhood aggregation. For DNA sequence alone (seq) and for DNA sequence combined with DNase (DNase), CHROME (Seq) and CHROME (DNase) were compared against CNN baselines following the EPCOT architecture. For Evo2 embeddings (Evo2) derived from the 7B sequence model, CHROME (Evo2) was compared against a matched MLP baseline. Perclass analyses revealed systematic AUPR improvements when non-random contacts were incorporated (Figure 2A). For visualization, Seq- and Evo2-based results were zoomed to the 0.3 AUPR range, but consistent gains were observed across all three input settings. Improvements were particularly evident in classes with more positive examples (larger yellow points). Seq- and Evo2-based models exhibited comparable performance ranges, indicating that although Evo2 embeddings are more compact, they do not surpass raw sequence features in this task—consistent with recent reports that foundation genomic models often fail to outperform simpler encodings [41]. In contrast, DNase-based models achieved substantially higher per-class scores, consistent with the added cell line-specific signal provided by accessibility profiles [42]. Cell line-specific scatter plots illustrating these results are shown in Figure S2.

**Figure 2:**
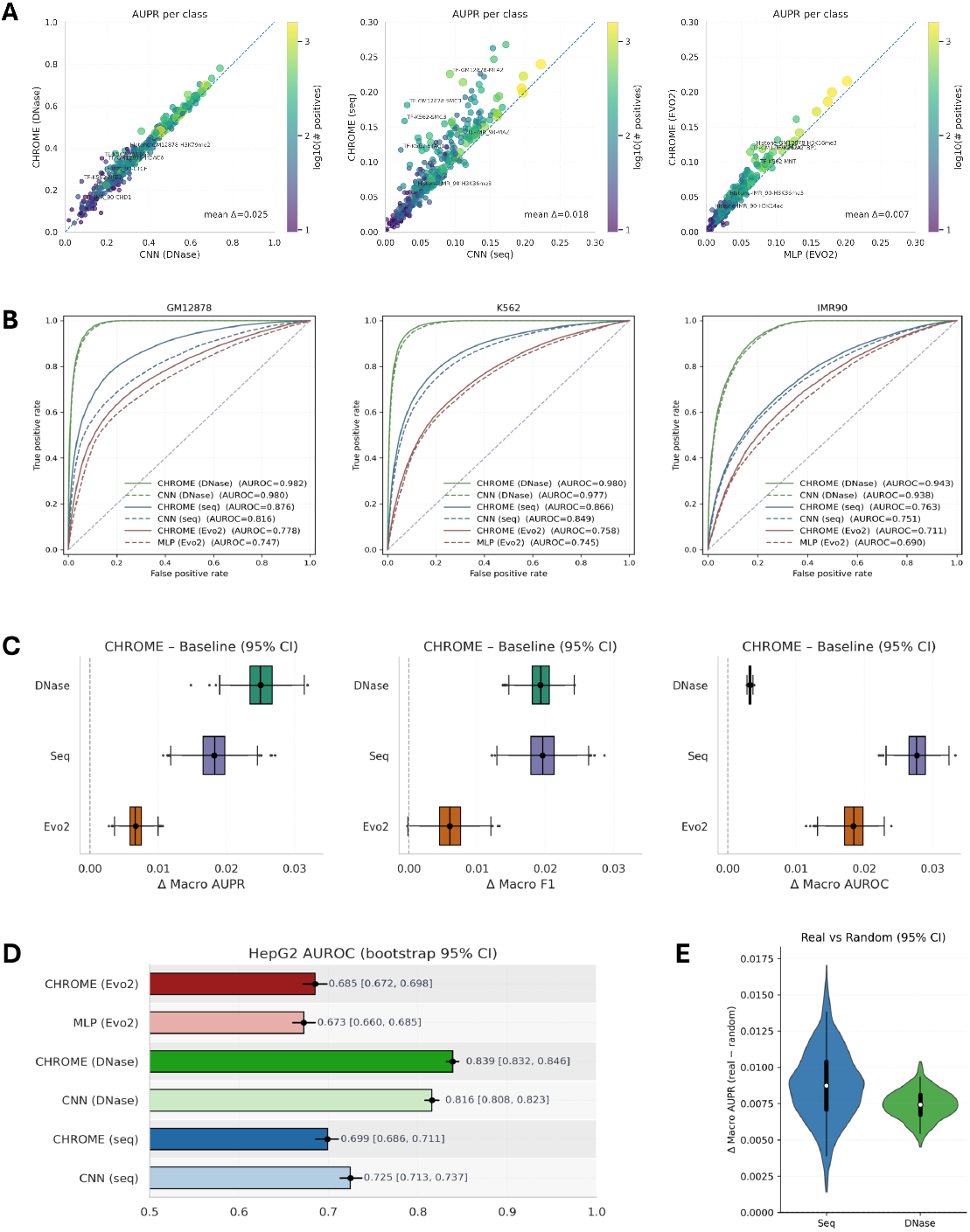
CHROME prediction of transcription factor and histone modification profiles. (**A**) Per-class AUPR compared with baselines. Left: DNase+DNA sequence (DNase); middle: DNA sequence only (Seq); right: Evo2 embeddings (Evo2). Each point corresponds to a transcription factor or histone modification, with color and size indicating the log_10_ number of positives. Analyses are restricted to classes with *>*30 positives. Seq and Evo2 results are zoomed to AUPR *≤* 0.30, while DNase shows the full range. The dotted line indicates *y* = *x*, and mean ΔAUPR is shown. (**B**) ROC curves for GM12878 (left), K562 (middle), and IMR-90 (right). CHROME (solid) shifts curves upward relative to baselines (dashed) across all input types. (**C**) Paired bootstrap estimates (95% CI) of CHROME *−* baseline differences in macro AUPR (left), macro F1 (middle), and macro AUROC (right), showing consistent positive shifts across feature sets. (**D**) Generalization to an unseen cell line (HepG2, chr9). Bars show AUROC with 95% bootstrap CIs. CHROME improves over baselines for DNase and Evo2, while Seq-only performs slightly below its baseline. (**E**) Ablation analysis with random controls. Real non-random contacts were replaced with degree- and distance-matched random Hi-C contacts (*±*100 kb). ΔMacro-AUPR (real *−* random) is positive for both Seq and DNase, confirming that non-random contacts provide more biologically meaningful signals.

ROC analyses across GM12878, K562, and IMR-90 revealed a consistent pattern (Figure 2B). For all three input types—DNase with Seq, Seq, and Evo2—CHROME shifted the ROC curves upward relative to their corresponding baselines. The magnitude of improvement scaled with input richness: DNase+Seq achieved the strongest overall performance, followed by Seq, while Evo2 embeddings performed worst in absolute terms. Nevertheless, in every setting the inclusion of non-random contacts yielded measurable gains.

Paired bootstrap analyses confirmed consistent improvements across input types, with distributions of Δ macro AUPR, F1, and AUROC all shifted above zero (Figure 2C). F1 scores were calculated using thresholds optimized on the validation set and then applied to the test set, ensuring consistent evaluation across models. Gains in AUPR were largest for DNase, followed by Seq and then Evo2. For F1, both DNase- and Seq-based models showed comparable improvements, with Evo2 again yielding smaller but positive shifts. In contrast, AUROC improvements were most pronounced for Seq, moderate for Evo2, and smallest for DNase, likely reflecting the already strong baseline performance of DNase. Together, these results indicate that CHROME enhances predictive performance across all feature types, with the magnitude of gain depending on both the metric and the underlying representation.

We also examined a cell line-specific training scenario, in which CHROME was trained, validated, and tested within a single cell line. Similar trends were observed, where incorporating non-random loci information improved signal prediction; however, the overall gain was smaller than that observed in cross-cell line training. For example, the IMR-90 CHROME (DNase) model showed a larger performance increase under cross-cell line training (Figure S5).

We next assessed the generalization capacity of CHROME in a cell line-independent setting by testing models trained on GM12878, K562, and IMR-90 against HepG2 profiles on chromosome 9. CHROME maintained strong performance and outperformed baselines when DNase or Evo2 embeddings were used (Figure 2D and Figure S3). In contrast, the sequence-only model performed slightly below its baseline, indicating that raw sequence features alone lack the cell line specificity required for cross-context transfer. By comparison, both DNase and Evo2 embeddings provided richer representations, capturing accessibility or dense sequence features that enabled CHROME’s neighborhood aggregation to yield systematic gains in an unseen cellular context. The single cell line-trained CHROME models were also tested on HepG2, showing transferable performance gains across input types, although the improvements were less stable and less significant than those observed in cross-cell line training (Figure S6).

Finally, we performed an ablation analysis on chromosome 9 for GM12878, IMR-90, and K562 to test whether CHROME’s gains depended on physically specific, non-random contacts. Each neighbor set was replaced with degree- and distance-matched random Hi-C loci sampled within 100 kb of each true non-random locus. Because DNase- and Seq-based models outperformed Evo2, we focused on these two settings. As shown in Figure 2E, graphs constructed from real non-random contacts consistently yielded higher macro AUPR than their randomized counterparts, with improvements reproducible across all three cell lines. The effect was slightly stronger for sequence models than for DNase, but significant in both. These results confirm that CHROME’s predictive advantage derives from biologically meaningful non-random contacts rather than from neighborhood size or linear genomic proximity alone.

### Contribution landscape of CHROME

Beyond predictive performance, CHROME provides mechanistic insight into how physically specific, non-random contacts influence regulatory activity. Using an attention mechanism [38, 43], it learns contribution weights that quantify each neighbor’s influence on the center locus, revealing how spatially connected loci shape predictions and highlighting the regulatory impact of specific chromatin interactions. As illustrated in Figure 3A, these learned weights can be ranked to assess neighbor-to-center influence, offering an interpretable framework for studying three-dimensional genome regulatory effects.

**Figure 3:**
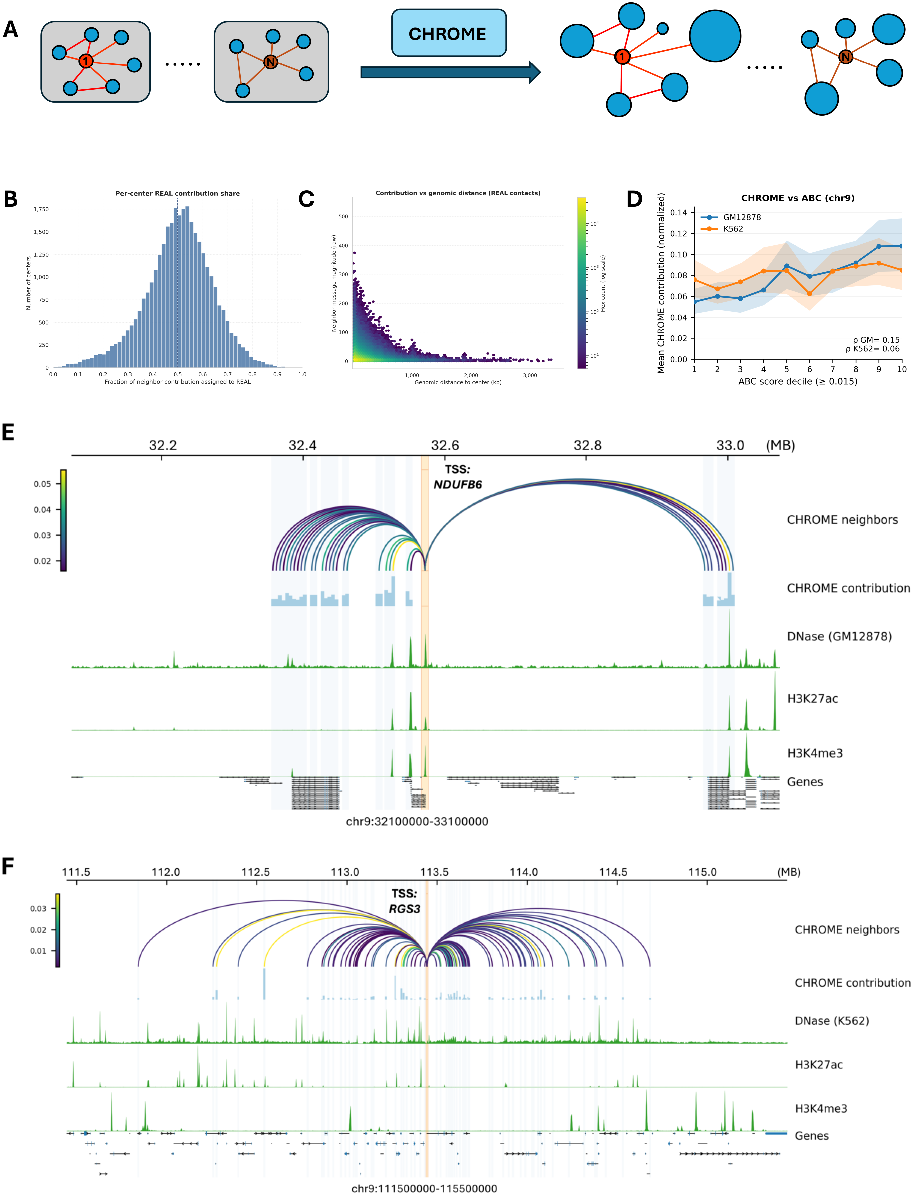
Contribution landscape learned by CHROME. (**A**) Schematic of the framework. Each 5 kb center locus is represented as a star graph with neighbors defined by non-random contacts. A graph attention model assigns edge-wise contributions from each neighbor to the center node, reflecting the relative influence of spatially connected loci on the center. **(B)** Distribution of the fraction of total neighbor-to-center contribution attributed to real non-random contacts compared with degree- and distance-matched random Hi-C controls under chr9. The distribution is skewed toward real non-random contacts, with the median above 0.5 (red dashed line), indicating preferential weighting of non-random interactions. **(C)** Relationship between neighbor-to-center contribution magnitude and genomic distance for real non-random contacts. Contributions decline within the first few hundred kilobases but remain consistently detectable at 1-2 Mb, with weaker yet measurable signals extending to 3-4 Mb. (**D**) Comparison with Activity-by-Contact (ABC) scores on chromosome 9. In both GM12878 (blue) and K562 (orange), CHROME (DNase) neighbor-to-center contributions increase across ABC score deciles, with moderate concordance observed (*ρ*_GM_ = 0.15, *ρ*_K562_ = 0.06), highlighting both agreement and divergence between the two approaches. (**E**) Local contribution map for NDUFB6 in GM12878. The TSS-centered locus is highlighted in orange, and non-random neighbors are shown as arcs colored by neighbor-to-center contribution strength. Bar plots indicate that several distal neighbors with strong contributions align with DNase hypersensitivity and active histone marks (H3K27ac, H3K4me3). (**F**) Long-range contribution map for RGS3 in K562. CHROME identifies multiple strong neighbor-to-center contributions spanning nearly 4 Mb, with arcs terminating in regions enriched for DNase and H3K27ac. These distal neighbors could represent potential regulatory loci influencing RGS3 through non-random three-dimensional chromatin contacts.

To test whether CHROME can distinguish contributions from true non-random contacts versus random Hi-C contacts, we extended the ablation dataset introduced in Figure 2E. Whereas the earlier setup replaced non-random contacts with random ones, here we combined non-random and degree- and distance-matched random Hi-C contacts, doubling the edge set. Using the trained CHROME (Dnase) model, we prioritized all contacts for each center locus on chromosome 9, pooling contribution scores from both real non-random and random neighbors. The resulting distribution of per-center contribution share was clearly skewed toward real non-random contacts, with the bulk of centers assigning more than half of the weight to real non-random neighbors (Figure 3B). This indicates that CHROME consistently prioritizes real non-random neighbors even when random Hi-C alternatives are present, showing that contribution weights capture biologically meaningful three-dimensional architecture rather than neighborhood size or linear proximity alone. As shown in Figure 3C, contributions from non-random contacts decay with distance but remain detectable at 1-2 Mb and extend weakly to 3-4 Mb.

To place CHROME in context with established enhancer-promoter frameworks, we compared CHROME (Dnase) neighbor-to-center contribution weights with Activity-by-Contact (ABC) scores [44], which combine enhancer activity with Hi-C contact frequency. ABC links, originally on hg19, were lifted to hg38 using UCSC liftOver [45] and then assigned to 5 kb loci by taking the maximum score per locus to match CHROME’s resolution. For each gene, we anchored on the TSS-centered 5 kb locus, collected its non-random Hi-C neighbors from the CHROME graph, and paired each neighbor’s normalized contribution with the corresponding binned ABC score. For analysis, we summarized the relationship by ABC-score deciles (threshold *≥* 0.015). On chromosome 9, CHROME contributions in GM12878 and K562 generally increased across ABC-score deciles but also showed discrepancies, with some loci receiving higher weight in CHROME despite modest ABC scores and vice versa (Figure 3D). While genome-wide correlations were not always aligned with ABC, CHROME nevertheless recapitulated many ABC-supported connections and highlighted additional neighbor-to-center relationships that may reflect regulatory mechanisms beyond activity-contact heuristics (Table S1).

We further visualized contribution landscapes for individual genes. For NDUFB6 in GM12878, CHROME (Dnase) highlighted a compact 1 Mb window around the TSS, with the TSS-centered 5 kb locus shaded in orange and all neighboring 5 kb loci shown in light blue (Figure 3E). Non-random neighbors were connected to the TSS by arcs, with arc color indicating contribution strength, and the same values were also represented as bar heights in a parallel track. Several neighbors with strong contributions coincided with DNase hypersensitivity and peaks of H3K27ac and H3K4me3, consistent with local regulatory activity. In contrast, for RGS3 in K562, CHROME revealed a broader landscape spanning nearly 4 Mb, again with neighbors shaded in light blue and contribution strength conveyed by both arc color and bar height (Figure 3F). Multiple distal neighbors contributed substantially to the TSS; some overlapped DNase and histone modification peaks, while others did not, suggesting a broader set of potential regulatory loci that may influence RGS3 through non-random contacts. Additional contribution visualizations for other representative loci are provided in Figures S7 and S8.

### Prediction and interpretation of tissue-specific eQTLs

We next evaluated whether the regulatory knowledge learned by CHROME from ChIP-seq profiles can be transferred to tissue-specific regulatory variation, using GTEx eQTLs [34] as an independent benchmark. To approximate relevant contexts, we mapped GM12878 to blood and IMR-90 to lung, reflecting their origins as lymphoblastoid cells and lung fibroblasts, respectively. CHROME models trained on ChIP-seq prediction were repurposed as embedding generators and applied to the same eQTL datasets curated in Borzoi [20], which include both positive and negative controls. For each variant, graph embeddings were generated for the variant-centered 5 kb locus using CHROME, and local sequence embeddings were generated for the TSS-centered 5 kb locus tested for association with the variant using the corresponding baseline model, for blood (GM12878) and lung (IMR-90). These embeddings were concatenated and passed to an MLP classifier to predict whether the variant corresponded to a true eQTL.

For comparison, CNN or MLP baselines were applied independently to both the variant- and TSS-centered 5 kb loci, after which their embeddings were concatenated and classified using the same MLP. This design ensured that CHROME- and baseline-derived embeddings were evaluated within an identical framework, allowing a direct test of whether the neighborhood context captured by CHROME provides a predictive signal beyond local-only encoders.

As shown in Figure 4A, precision-recall curves across five folds demonstrate that CHROME-based embeddings (solid lines) consistently outperform baseline embeddings (dashed lines). This advantage holds for both DNase- and sequence-based inputs, except Evo2 in lung, which performed slightly worse than its baseline. These trends are summarized in Figure 4B, where F1 scores with 95% confidence intervals confirm that CHROME (DNase) and CHROME (Seq) achieved the strongest overall performance, while Evo2 models were less robust. ROC curves and AUROC values across the same five folds showed consistent trends, as presented in Supplementary Figure S9A-C. Top-k precision and calibration analyses in blood and lung further support these results, showing that CHROME maintains higher precision among top-ranked variants and calibration comparable to or better than baselines (Supplementary Figure S9D-E). Together, these findings indicate that knowledge learned from non-random contacts in regulatory prediction transfers effectively to gene-variant association, yielding more accurate and interpretable eQTL classification than local-only baselines, particularly when cell line-specific features such as DNase are included.

**Figure 4:**
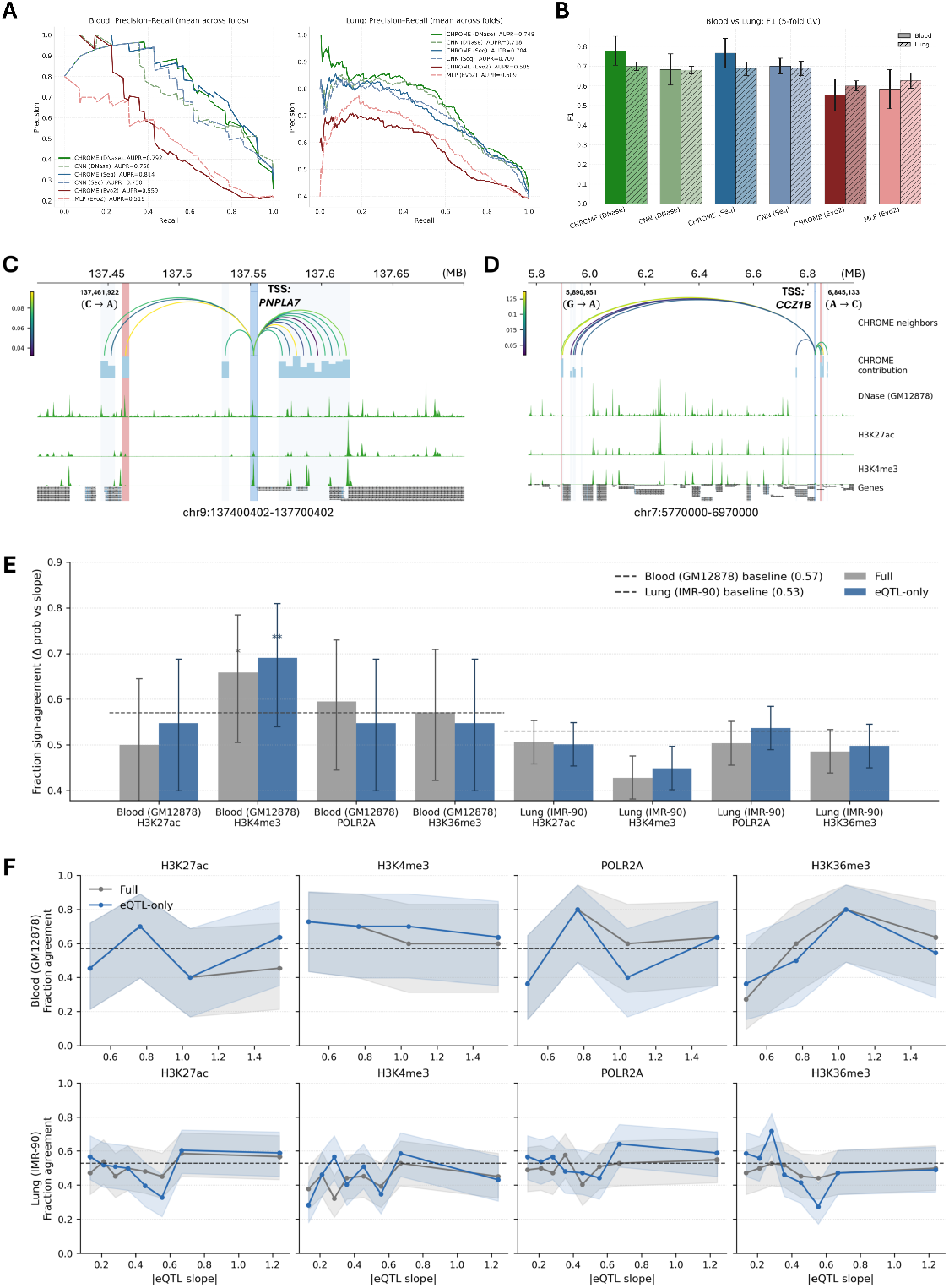
Evaluation of CHROME eQTL prediction and contribution interpretation. (**A**) Cross-validation performance in two representative GTEx tissues (blood and lung). Precision-recall curves show that CHROME embeddings (solid) outperform local baseline embeddings (dashed). (**B**) Summary bar plots (mean ± s.d. across five folds) of F1 scores in blood and lung, where CHROME models consistently achieve higher values than CNN baselines. (**C**) Contribution map for *PNPLA7* in blood (GM12878). The TSS-centered locus (light blue) is linked to a non-random neighbor containing the eQTL variant chr9:137,461,922 (C *→* A, red), with arcs colored by contribution strength and bar plots showing relative contributions. (**D**) Contribution map for *CCZ1B* in blood (GM12878). Strong contributions are observed from both a distal eQTL (chr7:5,890,951 G *→* A) and a local eQTL (chr7:6,845,133 A *→* C), despite limited overlap with DNase or histone peaks. (**E**) Aggregate sign-agreement analysis between CHROME (DNase)-predicted Δ probability and eQTL slopes for four representative marks in blood (GM12878) and lung (IMR-90). Bars show mean *±* 95% confidence intervals relative to baseline expectations derived from majority class proportions. (**F**) Stratified analysis of sign-agreement as a function of absolute eQTL effect size. Agreement fractions increase progressively in subsets with stronger eQTL slopes.

At the locus level, CHROME contribution maps reveal how distal neighbors mediate eQTL effects through their connections to the center TSS bin. Previous studies have shown that many-body chromatin interactions, such as those occurring in condensates, likely play important roles in gene regulation (46–49). Consistent with this, CHROME (DNase) highlights not only direct local contacts but also distal multi-locus interactions that can modulate promoter activity. In PNPLA7 (blood, Figure 4C), an eQTL variant at chr9:137,461,922 (C→A; red shading) lies within a non-random neighbor that contributes strongly to the TSS bin (blue shading). This neighbor coincides with DNase hypersensitivity and active histone marks, exemplifying a localized eQTL mediated by canonical regulatory signals. By contrast, in CCZ1B (blood, Figure 4D), both a distal eQTL (chr7:5,890,951 G→A) and a local eQTL (chr7:6,845,133 A→C) fall within neighbors that receive strong contribution weights to the TSS bin, despite lacking overlap with DNase or histone peaks. These examples demonstrate that CHROME can highlight both epigenetically supported neighbors and potential regulatory loci not readily explained by conventional chromatin features. In both cases, strong neighbor-to-TSS contributions provide a mechanistic link between genetic variation and promoter regulation. Additional examples in lung tissue, including KCNH1 and PCCA, are shown in Supplementary Figure S10.

Finally, we explored whether CHROME could provide insight into the directionality of eQTL effects, specifically whether a variant increases or decreases the expression of its target gene. This remains a difficult challenge: most approaches fail due to limited receptive fields or reliance on a reference genome (14, 16, 50–52), and even with personalized genomes [53], the problem has not been solved. Although CHROME was not trained to predict quantitative expression changes, its contribution framework enables comparison between predicted shifts in gene expression-associated ChIP-seq profiles and GTEx eQTL slopes. We focused on four representative marks—H3K27ac, H3K4me3, POLR2A, and H3K36me3—which are all positively related to transcription.

Two settings were evaluated: one including only the eQTL-harboring neighbor to enforce a one-to-one relationship with the TSS, and another embedding the eQTL within the full set of non-random neighbors. Sign-agreement between CHROME (DNase)-predicted Δ probabilities (mut vs ref) and GTEx slopes was then measured, with the majority proportion of up-vs. down-regulating variants serving as baseline. In blood (GM12878), H3K4me3 showed agreement clearly above baseline in both settings, while POLR2A surpassed baseline only when the full neighborhood was included (Figure 4E). In lung (IMR-90), overall agreement did not exceed baseline across marks, but stratified analyses revealed a consistent trend: as the absolute eQTL slope increased, CHROME’s sign-agreement also rose. For H3K27ac and POLR2A in particular, high-slope subsets showed stronger agreement beyond baseline expectations (Figure 4F). Together, even indirectly, CHROME provides greater predictive power for eQTL directionality than models that treat DNA as a purely sequential input, by incorporating non-random chromatin contacts that integrate regulatory information from both cis and distal loci. Directional agreement is strongest at variants with pronounced regulatory impact, and including the full non-random neighborhood can enhance concordance in select cases, such as POLR2A in blood.

### Variant pathogenicity prediction and non-coding regulatory disruption

We next tested whether CHROME’s regulatory knowledge, derived from non-random chromatin contacts, can be transferred to distinguish pathogenic from benign variants. For this benchmark, we used curated ClinVar [35] datasets containing single-nucleotide variants and short indels annotated as benign or pathogenic. Because ClinVar does not provide cell line- or tissue-specific information, we constructed a consensus input representation in which DNase accessibility was averaged across GM12878, K562, and IMR-90 to approximate a general open chromatin state, and non-random Hi-C contacts were defined as the intersection across these three cell lines to represent the most common contact patterns. Each variant was placed at the center of its 5 kb locus, and graph embeddings were generated by aggregating information from these consensus non-random contacts using trained CHROME models. In parallel, matched baseline models (CNN or MLP) produced embeddings from the variant locus in isolation. All embeddings were then passed to an MLP classifier to predict variant pathogenicity, directly testing whether CHROME’s contextual information improves upon local-only encoders.

Across five-fold validation, CHROME-based embedding consistently outperformed their baseline counterparts (Figure 5A-C). Precision-recall curves showed the greatest gains for sequence- and Evo2-based inputs, while calibration analyses indicated that CHROME-derived probabilities aligned more closely with empirical frequencies, resulting in lower or comparable Brier scores and expected calibration errors. Violin plots further confirmed a clearer separation between benign and pathogenic variants under CHROME than baselines. DNase-based embeddings, by contrast, performed substantially worse than sequence-or Evo2-based embeddings regardless of whether neighborhood context was used. This likely reflects the limitations of averaging DNase accessibility across cell lines, which fails to capture canonical regulatory activity patterns [39, 54]. Nevertheless, adding the consensus set of non-random contacts improved performance across all feature types, including DNase, demonstrating that non-random chromatin contacts convey a robust and transferable predictive signal, even when input features are less informative. Additional performance analyses, including ROC curves, top-k precision, cumulative gain, and operating-point evaluations, are provided in Supplementary Figure S11, further corroborating CHROME’s superior predictive and clinical relevance.

**Figure 5:**
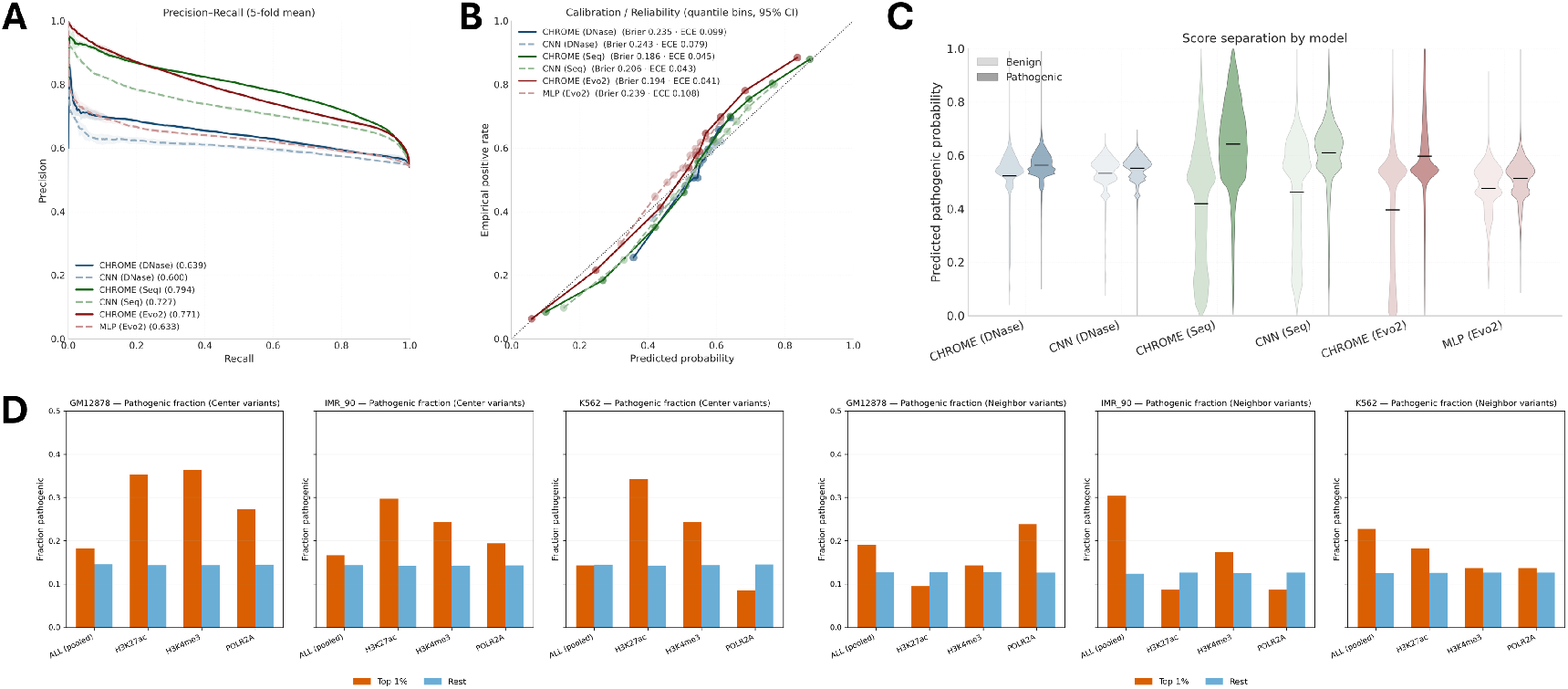
ClinVar variant classification and non-coding pathogenicity. (A) Precision-recall curves (5-fold mean) for MLP classifiers trained on CHROME graph-based embeddings versus baseline variant-locus embeddings. CHROME consistently outperforms CNN and MLP baselines, with the largest gains observed for sequence- and Evo2-based inputs. (B) Calibration plots comparing predicted probabilities with empirical frequencies. All three CHROME variants (Seq, Evo2, DNase) show improved calibration over their baselines, reflected in lower or comparable Brier scores and expected calibration error. (C) Violin plots of predicted pathogenic probabilities for benign versus pathogenic mutations. CHROME achieves clearer class separation than baselines, particularly for sequence and Evo2 inputs. (D) Pathogenic enrichment stratified by absolute probability change (| Δprob—). Left: cis mutations located in the variant-centered bin. Right: distal mutations occurring in non-random neighbors. In all cell lines, the top 1% most perturbed mutations are enriched for pathogenic cases relative to the remainder, with pooled analyses showing especially strong enrichment for distal mutations.

To directly assess non-coding pathogenicity, we used ncVarDB [55], a manually curated collection of pathogenic non-coding mutations paired with benign controls. We use CHROME (DNase) to evaluate whether predicted regulatory disruptions correlate with pathogenic status. For each mutation, we generated predictions for both reference and mutant alleles across cell line-specific ChIP-seq profiles and quantified the absolute probability change (|Δprob|) when the mutation occurred in either the center locus or a non-random neighbor locus. Mutations were then stratified by |Δprob|, contrasting the top 1% most perturbed against the remainder. In this analysis (Figure 5D), pathogenic enrichment was evident for both cis mutations located in the center locus and distal mutations occurring in non-random neighbor loci. Although both categories showed clear enrichment within the top 1%, the effect was consistently stronger for distal mutations when results were pooled across profiles. This pattern indicates that pathogenic non-coding variants can disrupt regulatory activity through both local and distal mechanisms, with distal perturbations showing especially pronounced effects in aggregate.

## DISCUSSION

Our goal was to develop a predictive genomics framework that leverages three-dimensional chromatin architecture to model regulatory interactions across both cis and distal loci. Gene regulation depends on coordinated interactions among DNA sequence, chromatin state, and spatial organization, yet most existing models still treat the genome as linear, overlooking its three-dimensional context. Although Hi-C provides a global view of chromatin folding, its contact maps are population-averaged, distance-biased, and noisy, making biologically specific interactions difficult to extract and integrate into predictive models. To overcome these challenges, we used CHROMATIX [33], which physically validates Hi-C data via a self-avoiding polymer ensemble null model, filtering out random noise and retaining only physically specific, non-random contacts. Recent studies have shown that integrating such physically constrained interactions enhances the mapping of non-coding variants to their regulatory targets [56, 57]. These non-random contacts form the foundation of CHROME, a chromatin-structure-guided graph attention framework that models information flow through biologically meaningful spatial connections. The framework extends the receptive field to 4 Mb, supports flexible sequence and epigenetic embeddings, and provides intrinsic interpretability by assigning attention-based contribution weights from each neighbor to the center locus.

On the benchmark task of predicting transcription factor and histone modification ChIP-seq profiles, CHROME consistently outperformed local encoder-only baselines across GM12878, K562, and IMR-90, and generalized robustly to the unseen HepG2 cell line. Incorporating non-random contacts improved predictive performance across all input types, including raw sequence, DNase accessibility, and Evo2 embeddings, demonstrating that chromatin structure complements diverse feature embeddings by introducing spatial context otherwise absent from linear models. Ablation analyses confirmed that non-random contacts outperform degree- and distance-matched random Hi-C contacts, indicating that CHROME’s advantage arises from biologically meaningful spatial specificity rather than graph size or density.

Beyond regulatory prediction, embeddings derived from trained CHROME models improved tissue-specific eQTL classification and ClinVar variant pathogenicity prediction. Mapping GM12878 to blood and IMR-90 to lung tissues from GTEx [34, 20], CHROME embeddings consistently outperformed local-only baselines in eQTL prediction. Contribution maps revealed both canonical high-impact eQTL loci coinciding with DNase accessibility and histone marks, and non-canonical loci supported primarily by non-random contacts. Analyses of sign agreement with GTEx slopes further showed that CHROME can partially capture eQTL directionality by comparing predicted expression-related ChIP-seq changes between mutant and wild-type alleles, with the strongest concordance observed for variants with large effect sizes. Extending to clinical variation, CHROME graph embeddings also improved ClinVar pathogenicity prediction [35], outperforming CNN and MLP baselines. Non-coding pathogenic variants were enriched in both cis and distal loci that showed large predicted regulatory perturbations, consistent with previous findings that disease-associated mutations often occur in regulatory elements [58, 59], while further demonstrating how non-random chromatin contacts amplify these functional effects.

Our findings establish that physically validated non-random contacts are not merely structural but functionally informative for gene regulation. Incorporating these contacts improves predictive accuracy across all input modalities and reveals functional spatial relationships that linear sequence models cannot capture. Moreover, CHROME’s attention-based framework provides mechanistic interpretability, showing how individual contacts, both proximal and multi-megabase apart, influence regulatory activity. Contribution landscapes revealed that neighbor-to-center influences decay with distance but remain detectable at 1 to 2 MB, with weaker signals still measurable at 3 to 4 MB. These long-range contributions are consistent with reports of developmental enhancers acting across megabase distances and enhancer-promoter compatibility spanning multi-megabase loci [60, 61]. Together, these insights bridge predictive modeling with biological mechanism, providing a unified view of how three-dimensional chromatin organization influences regulatory function.

Conceptually, CHROME bridges physics-based modeling, graph learning, and foundation scale genomics. By coupling physically validated non-random contacts with a graph attention architecture, it integrates the mechanistic principles of genome folding with data-driven representation learning. This physics-informed design complements sequence-centric frameworks such as Enformer [16] and multimodal architectures such as EPCOT [18], which extend receptive fields through self-attention across long linear windows but lack explicit spatial topology. Likewise, CHROME complements large pretrained sequence models such as Evo2 [19] and AlphaGenome [22], which capture molecular scale sequence features but remain blind to higher-order chromatin organization. By modeling regulatory geometry through physically constrained graph representations, CHROME provides interpretability grounded in three-dimensional genome topology and offers a pathway for integrating sequence embeddings with spatial context to achieve multiscale genomic prediction. Biologically, CHROME highlights how specific three-dimensional contacts modulate transcriptional activity, yielding interpretable hypotheses for enhancer-promoter compatibility and long-range regulation. By linking chromatin structure to functional consequence, CHROME establishes a foundation for structure-aware regulatory modeling that can extend to larger, multimodal, and single-cell contexts.

Currently, CHROME leverages high-quality Hi-C data from three cell lines, yet already demonstrates robust performance across tasks. As more deeply sequenced Hi-C datasets become available across tissues [62], developmental stages, and disease contexts, CHROME can evolve into a more comprehensive framework incorporating broader cell-type-specific and dynamic contacts. The receptive field, currently extended to 4 Mb, could be further expanded since some TADs span tens of megabases, and deeper or hierarchical graph architectures may incorporate additional layers of regulatory information derived from non-random contact maps. Because the current implementation relies on bulk Hi-C data that averages across heterogeneous cell populations, applying CHROME to single-cell, allele-specific, or temporal Hi-C datasets [63, 64] may reveal dynamic and cell-state-dependent regulatory interactions. By uniting predictive modeling with mechanistic interpretation, CHROME establishes a scalable foundation for chromatin structure-aware genomics that links three-dimensional genome architecture to functional regulation.

## DATA AVAILABILITY

All genomic data used in CHROME were aligned to the hg38 reference genome. DNase-seq and ChIP-seq profiles were downloaded from ENCODE [39], while eQTL datasets were obtained from GTEx v8 [34, 20]. Hi-C contact maps for GM12878, K562, IMR-90, and HepG2 were downloaded from the 4D Nucleome Consortium [9, 36]. Variant datasets were obtained from ClinVar [35] and ncVarDB [55].

Source code implementing CHROME, including training, and evaluation pipelines, is provided at GitHub (https://github.com/boweiye2u/CHROME).

## 4 FUNDING

The grant support of NIH R35GM127084 is gratefully acknowledged.

### 4.0.1 Conflict of interest statement

None declared.

## Notes

### Competing Interest Statement

The authors have declared no competing interest.

