## supplemental info for "A Chromatin-Structure-Guided Framework for Predictive and Interpretable Regulatory Genomics"

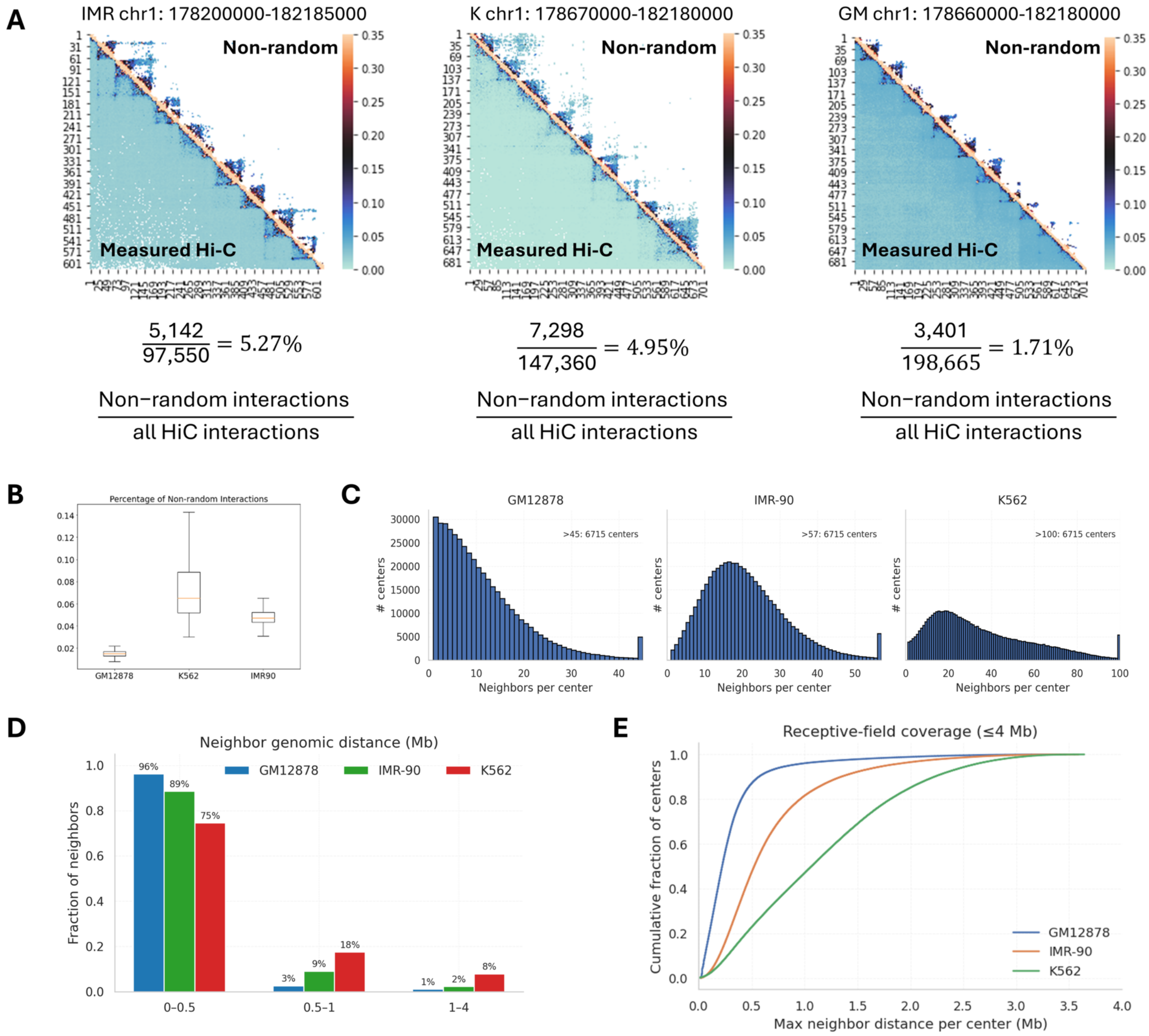


**Supplementary Figure S1: Hi-C preprocessing and graph statistics.** **(A)** Representative 4 Mb regions from IMR-90, K562, and GM12878 showing measured Hi-C contact matrices (bottom-left) and corresponding non-random contacts (top-right) identified by CHROMATIX. The ratio of non-random to total Hi-C contacts is shown for each region. **(B)** Genome-wide distribution of non-random interaction fractions across all 4 Mb regions for GM12878, IMR-90, and K562. **(C)** Degree distributions of CHROME graphs, showing the number of non-random neighbors per 5 kb center locus for each cell line. **(D)** Fraction of neighbors stratified by genomic distance loci (0–0.5, 0.5–1, 1–4 Mb), illustrating that most neighbors occur within sub-megabase distances but substantial long-range contacts are retained. **(E)** Receptive-field coverage quantified as the cumulative distribution of maximum neighbor distance per center locus, capped at 4 Mb. GM12878 reaches saturation more quickly, whereas IMR-90 and K562 show broader coverage across multi-megabase ranges.

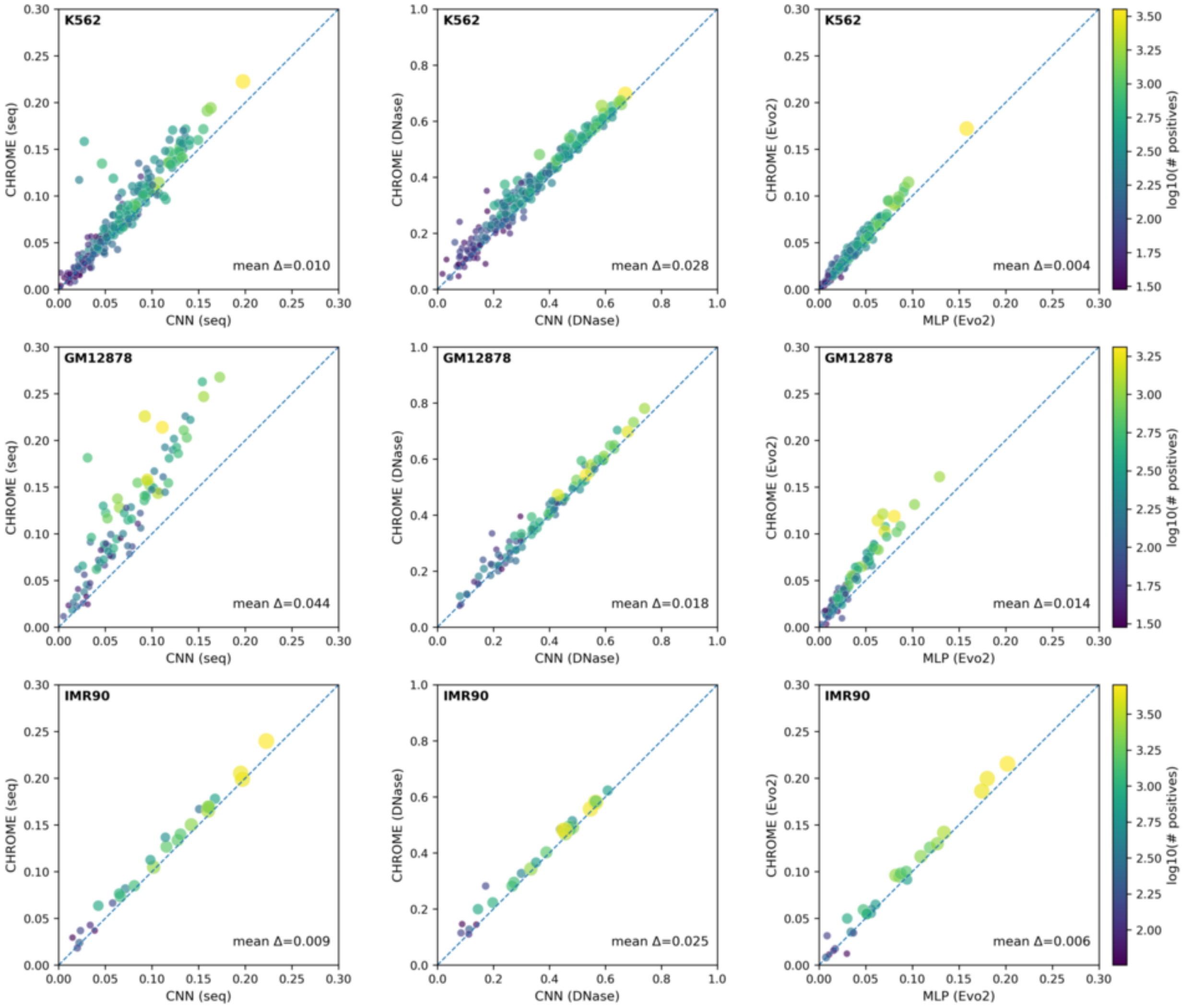


**Supplementary Figure S2: Per-class AUPR comparisons between CHROME and baseline encoders across cell lines.** Scatter plots show transcription factor and histone modification classes in K562 (top row), GM12878 (middle row), and IMR-90 (bottom row). Each point represents a class, colored by the $\log_{10}$ number of positive samples. The $x$-axis shows baseline performance (CNN for Seq and DNase inputs, MLP for Evo2 embeddings), and the $y$-axis shows CHROME performance with the same input features. The diagonal dashed line indicates parity between CHROME and baseline, and mean ΔAUPR values are reported in each panel. Analyses are restricted to classes with $\geq$30 positives. CHROME generally shifts points above the diagonal, indicating consistent performance gains across feature types and cell lines.

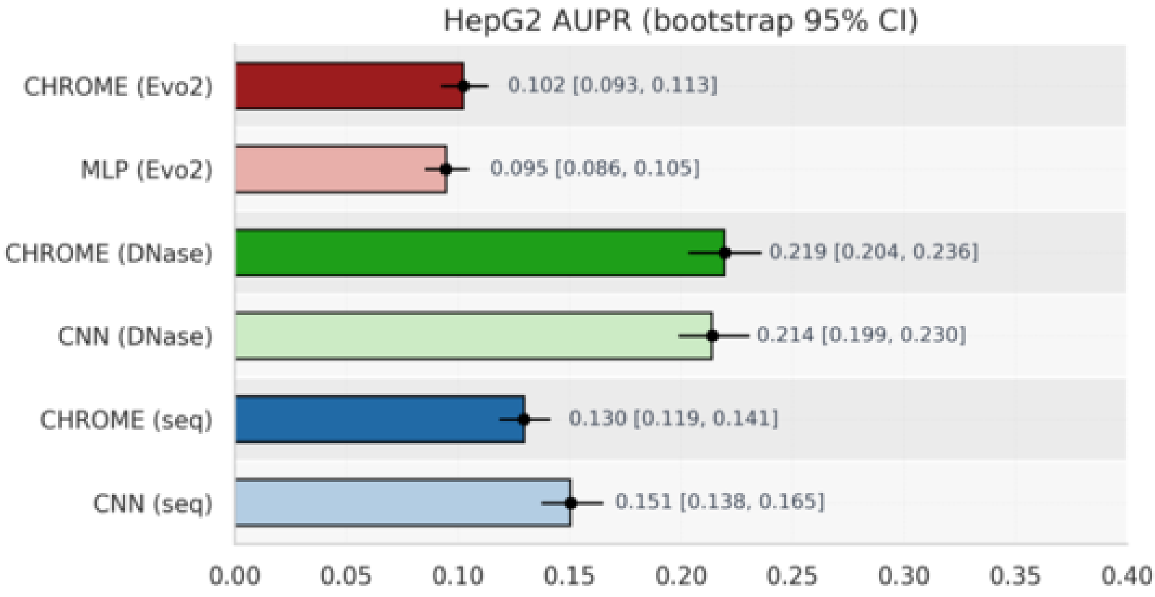


**Supplementary Figure S3: HepG2 performance on an unseen cell line.** Area under the precision–recall curve (AUPR) with 95% bootstrap confidence intervals for CHROME and corresponding baseline models on HepG2 chromosome 9. CHROME exhibits the strongest performance for DNase-based inputs and higher AUPR for Evo2 embeddings compared to the baseline, while its sequence-only variant performs marginally below the CNN baseline.

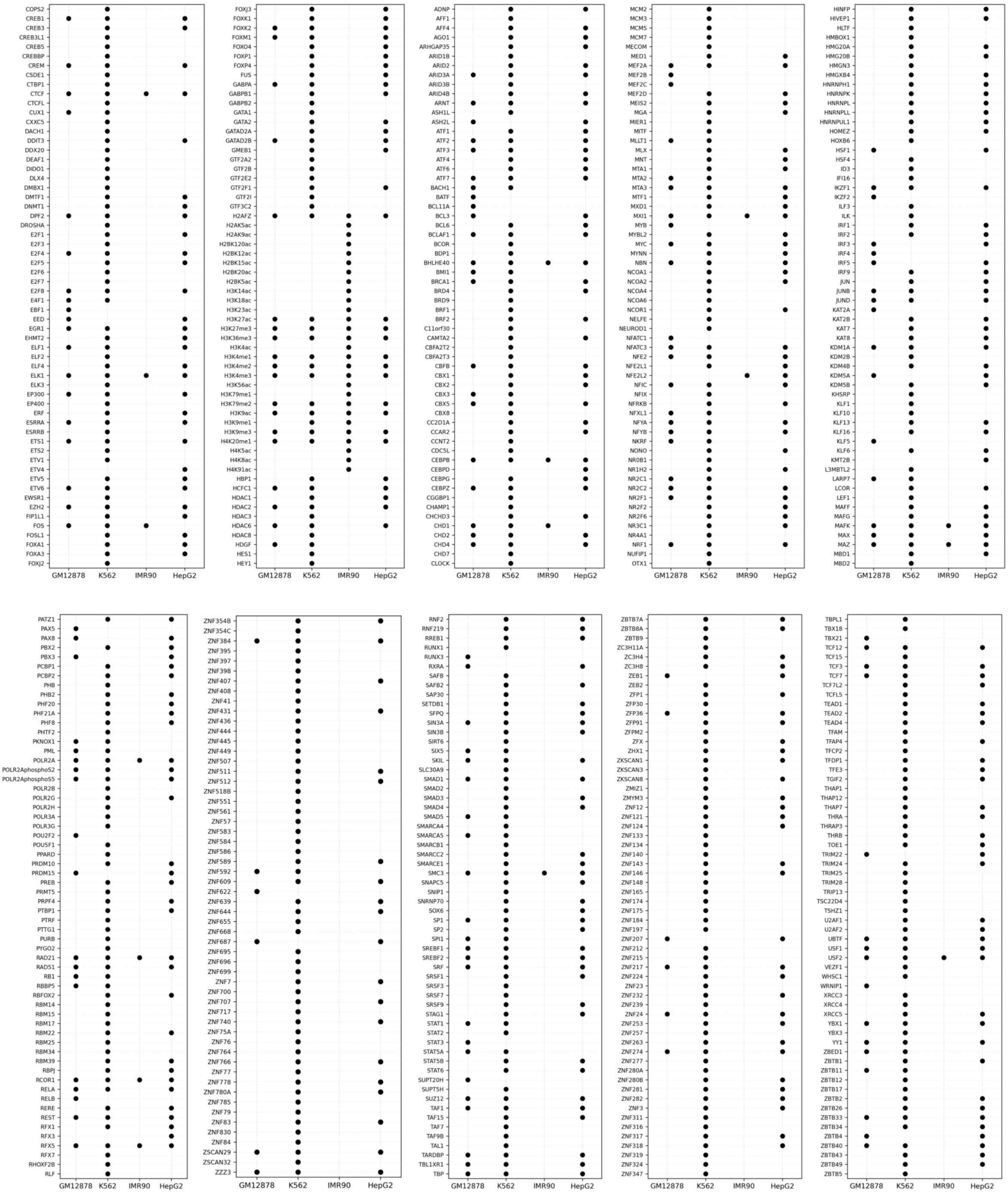

**Supplementary Figure S4: ChIP-seq availability across four cell lines.** Presence/absence table of transcription factor and histone modification ChIP-seq profiles collected from ENCODE [1] for GM12878 (168), K562 (539), IMR-90 (44), and HepG2 (324). Rows correspond to individual ChIP-seq targets, and filled circles indicate the availability of a narrowPeak dataset in the corresponding cell line.

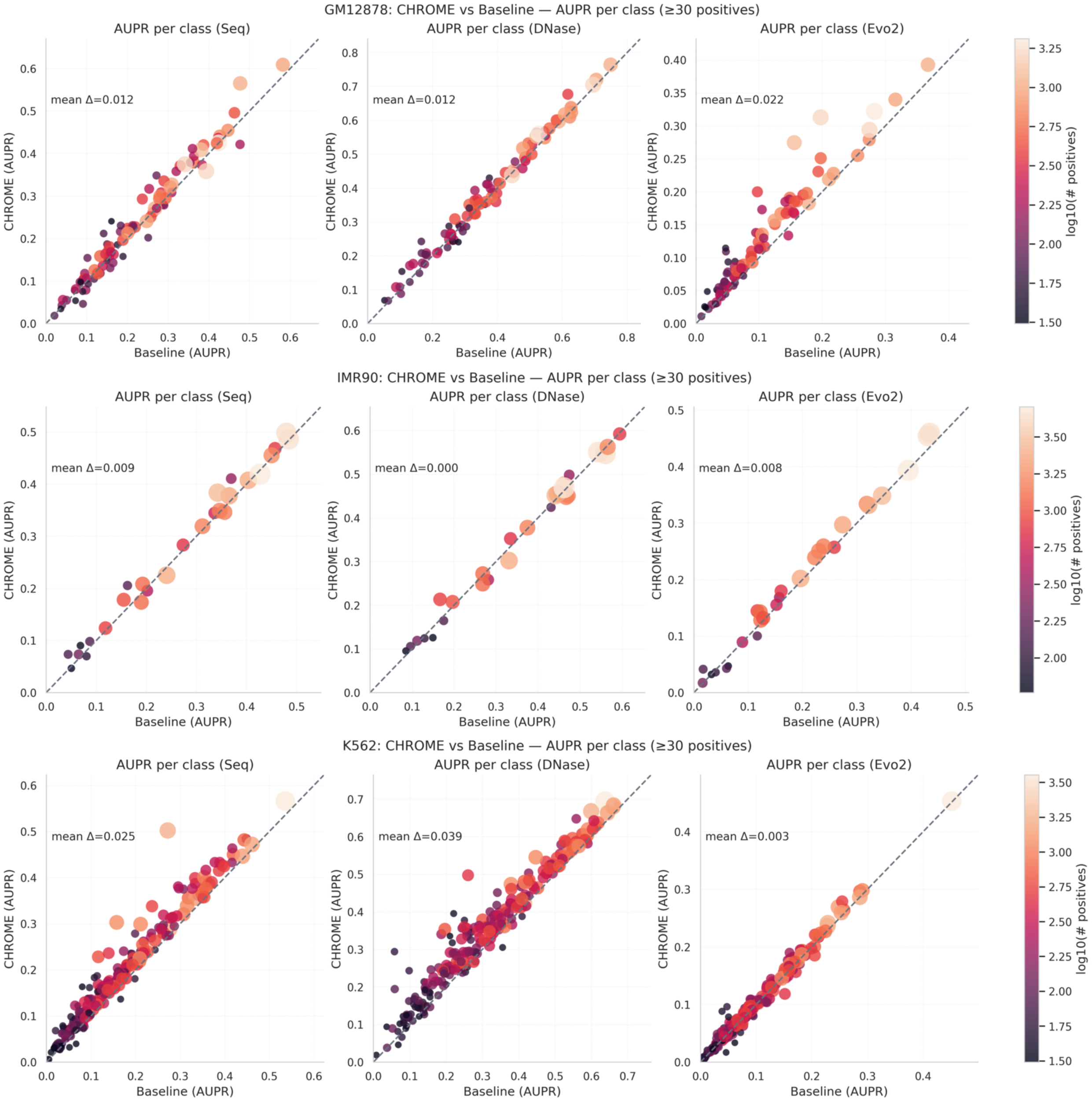


**Supplementary Figure S5: Cell line–specific training improves per-class prediction performance.** Scatter plots show per-class AUPR of CHROME compared to baseline models when trained and evaluated within the same cell line: (Top) GM12878, (Middle) IMR-90, and (Bottom) K562. Each dot represents a ChIP-seq profile ($\geq$30 positive loci), with dot size and color reflecting the number of positives ($\log_{10}$ scale). The diagonal line indicates parity with the baseline, and the mean $\Delta$ (average AUPR improvement of CHROME over the baseline) is reported in each panel. CHROME consistently achieves robust gains over baselines across sequence, DNase, and Evo2 feature settings in a cell line–specific manner.

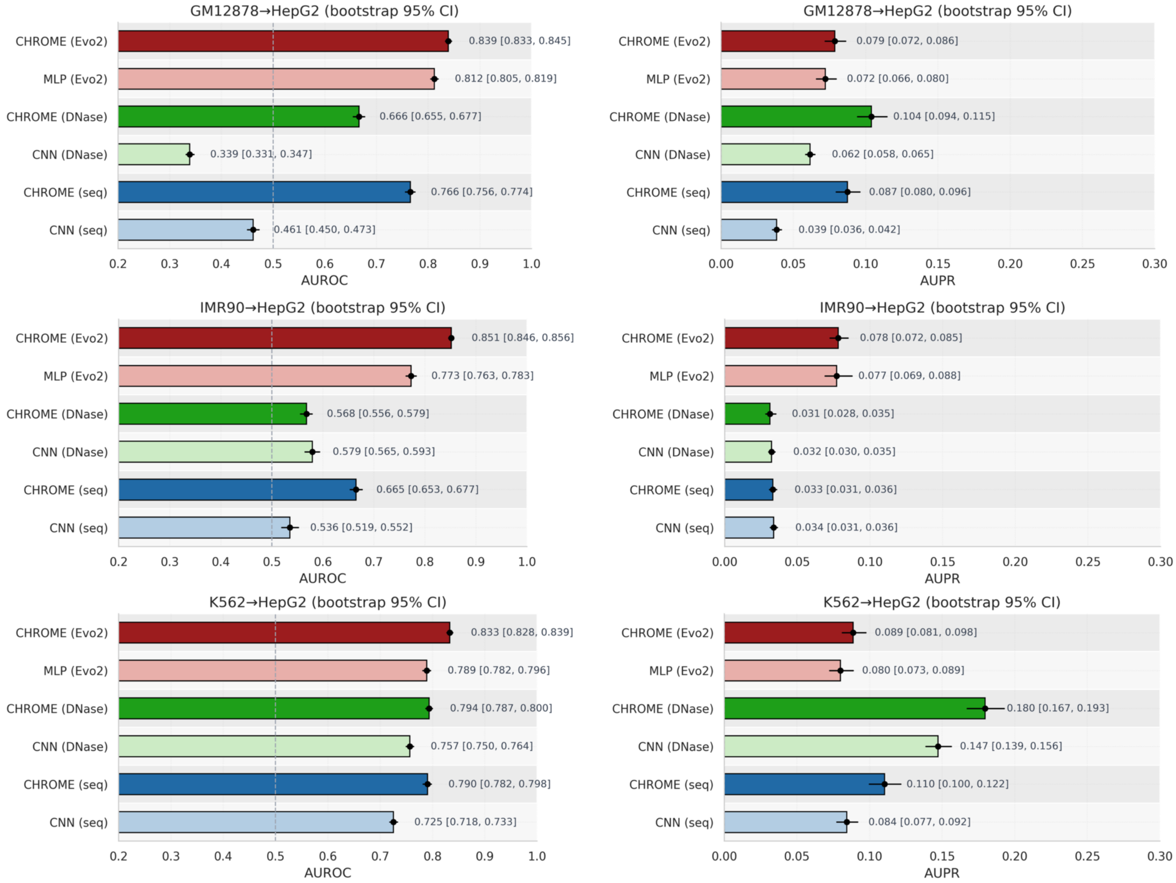


**Supplementary Figure S6: Cross-cell line generalization of single cell line–based CHROME models to unseen HepG2 chromosome 9 data.** Models were trained on (Top) GM12878, (Middle) IMR-90, or (Bottom) K562 ChIP-seq profiles and evaluated on the subset of HepG2 profiles shared with each training set. Left panels: AUROC with 95% bootstrap confidence intervals. Right panels: AUPR with 95% bootstrap confidence intervals, computed for ChIP-seq classes with at least 30 positive samples ($\geq 30$). Across all settings, CHROME consistently outperforms matched CNN or MLP baselines, with the strongest gains observed for Evo2-informed models (red) and DNase-informed models (green). The IMR-90–trained model shows lower overall performance primarily due to the limited number of overlapping ChIP-seq profiles available for evaluation. These results demonstrate that CHROME's neighborhood-aware framework generalizes beyond the training cell line and captures transferable regulatory signals even on an unseen chromosome.

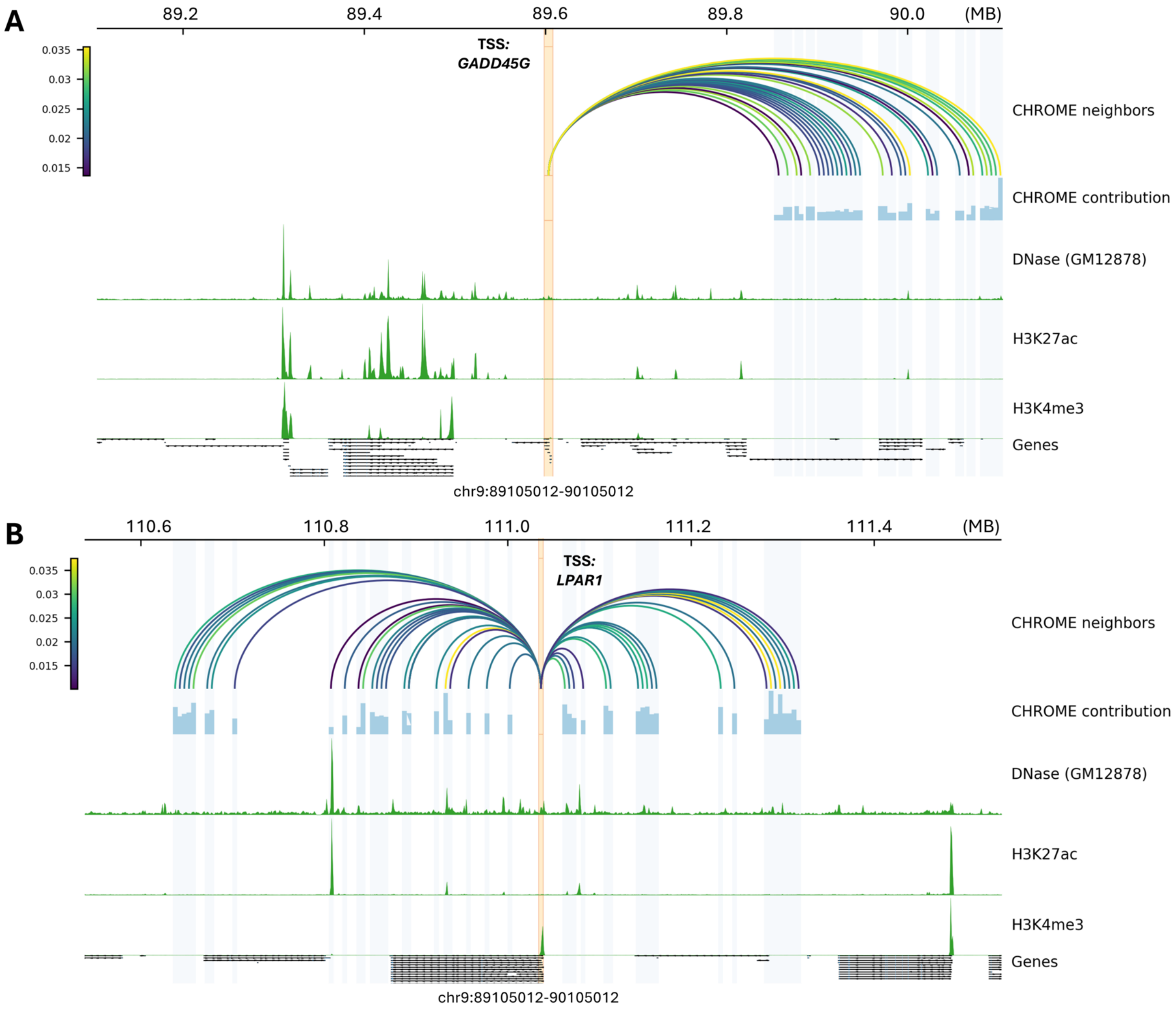


**Supplementary Figure S7: CHROME contribution profiles for GM12878.** (A) Spatially connected non-random Hi-C loci contributing to the 5 kb TSS-centered locus of *GADD45G* within a ∼1 Mb window. (B) Similar contributions to the 5 kb TSS-centered locus of *LPAR1* within a ∼1 Mb window. Arcs denote non-random Hi-C neighbors color-coded by contribution magnitude, while lower tracks show CHROME contribution scores, DNase accessibility (GM12878), histone modifications (H3K27ac, H3K4me3), and gene models.

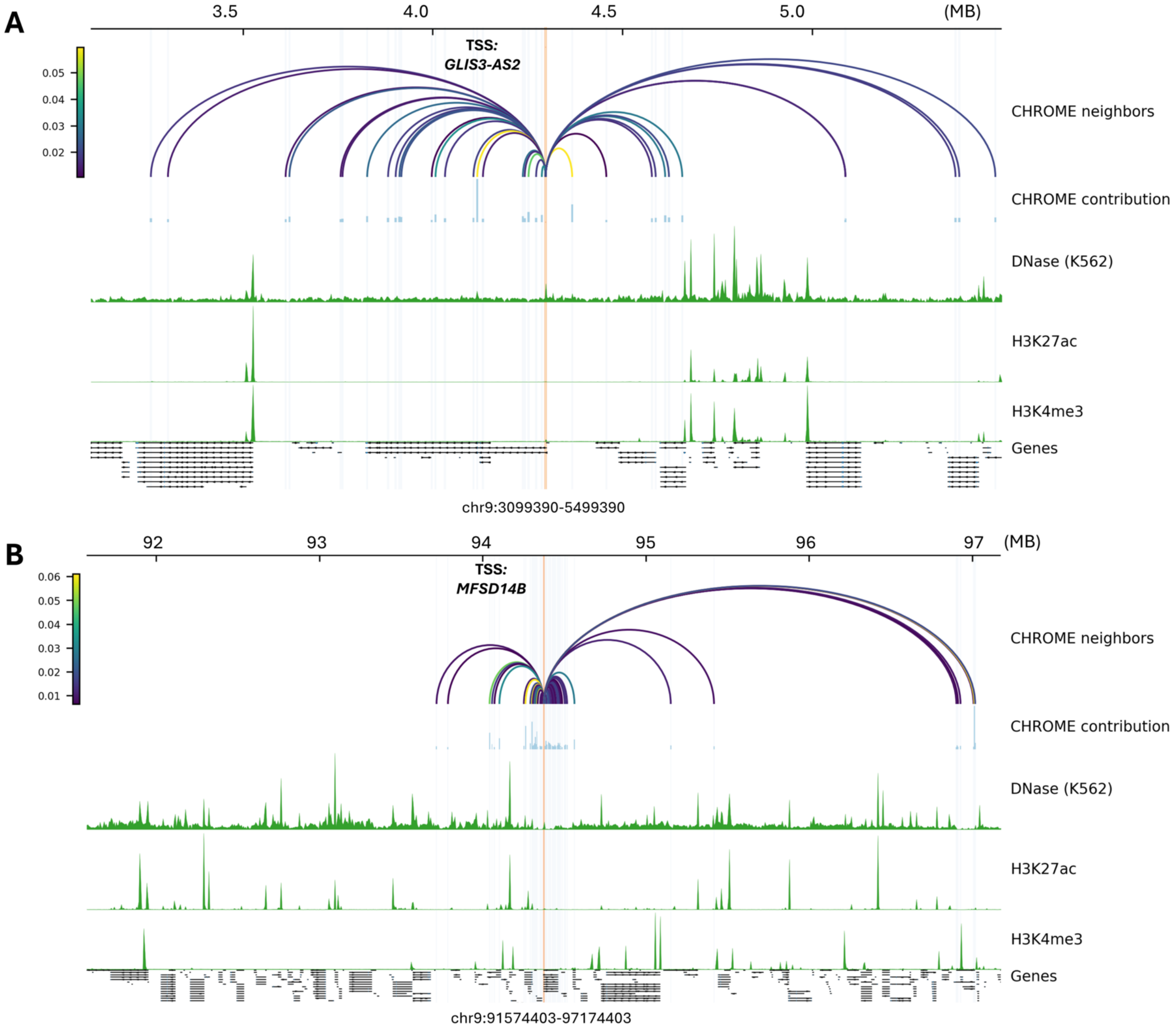


**Supplementary Figure S8: CHROME contribution maps reveal long-range regulatory effects in the K562 cell line.** **(A)** At the *GLIS3-AS2* locus, CHROME identifies spatially connected non-random Hi-C neighbors spanning approximately 2.5 Mb around the TSS-centered 5 kb locus. **(B)** At the *MFSD14B* locus, CHROME captures both a compact local regulatory neighborhood within ∼1 Mb of the TSS-centered locus and additional distal regulatory neighbors extending up to 3 Mb away, with dominant contributions concentrated near the proximal region. Together, these examples demonstrate that CHROME detects both proximal and distal regulatory influences in K562, with variable spatial ranges depending on gene context. Tracks shown include CHROME neighbor arcs, CHROME contribution scores, DNase accessibility, histone modifications (H3K27ac, H3K4me3), and gene models.

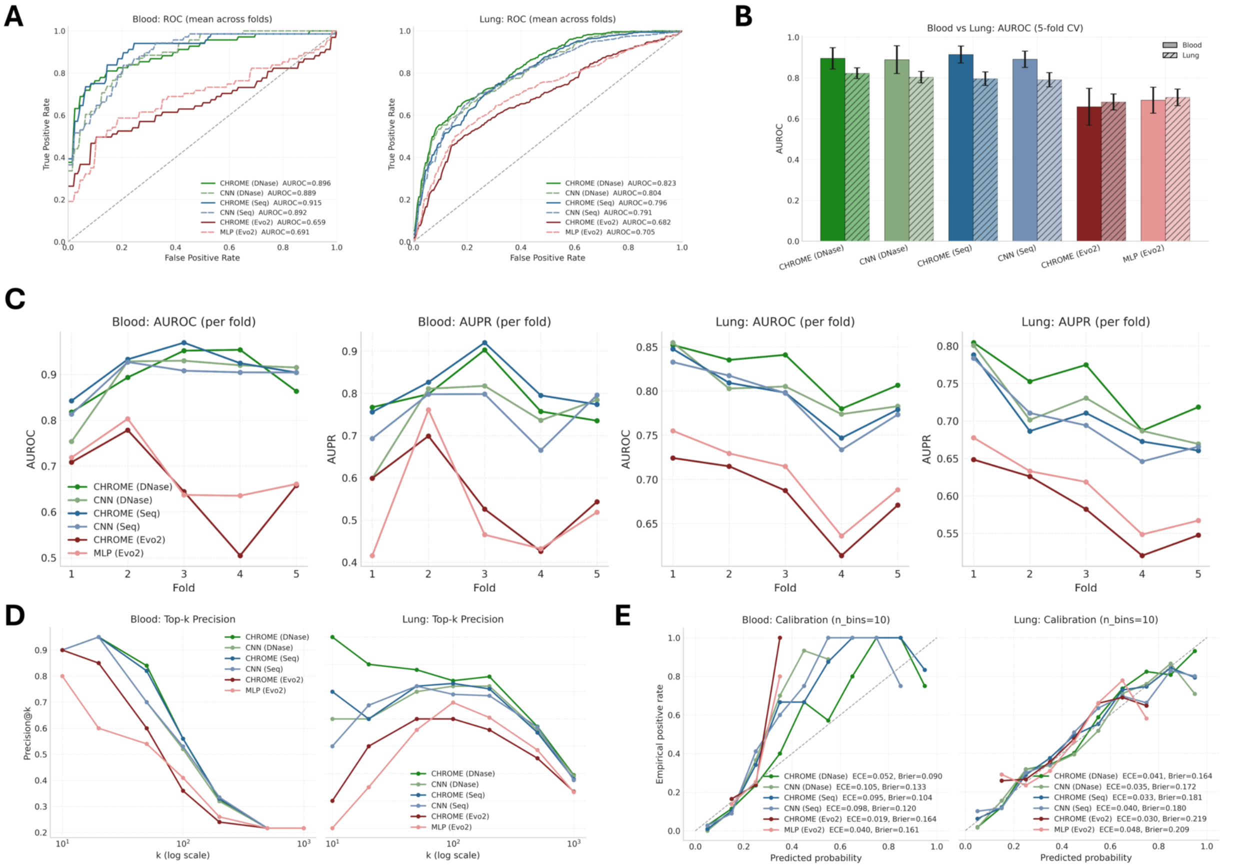


**Supplementary Figure S9: Supplementary evaluation of CHROME for eQTL classification across tissues.** **(A)** ROC curves (5-fold mean) for Blood and Lung tissues, with corresponding AUROC values reported per model. **(B)** Cross-tissue comparison of AUROC (5-fold mean ± 95% CI), showing consistently higher performance for GAT models relative to CNN baselines while the Evo2 version is slightly worsen than the baseline. **(C)** Per-fold AUROC and AUPR values in Blood and Lung. **(D)** Top-$k$ precision analysis in Blood and Lung (log-scaled $k$), indicating that CHROME maintains higher precision among top-ranked variants compared to baselines. **(E)** Calibration plots (10 bins) for Blood and Lung, reporting expected calibration error (ECE) and Brier score per model. Overall, CHROME (Seq, DNase) exhibits calibration that is comparable or favorable relative to the corresponding CNN baselines. Together, these analyses corroborate the main findings: CHROME improves AUROC/AUPR, maintains competitive calibration, and more effectively prioritizes top-ranked variants across tissues.

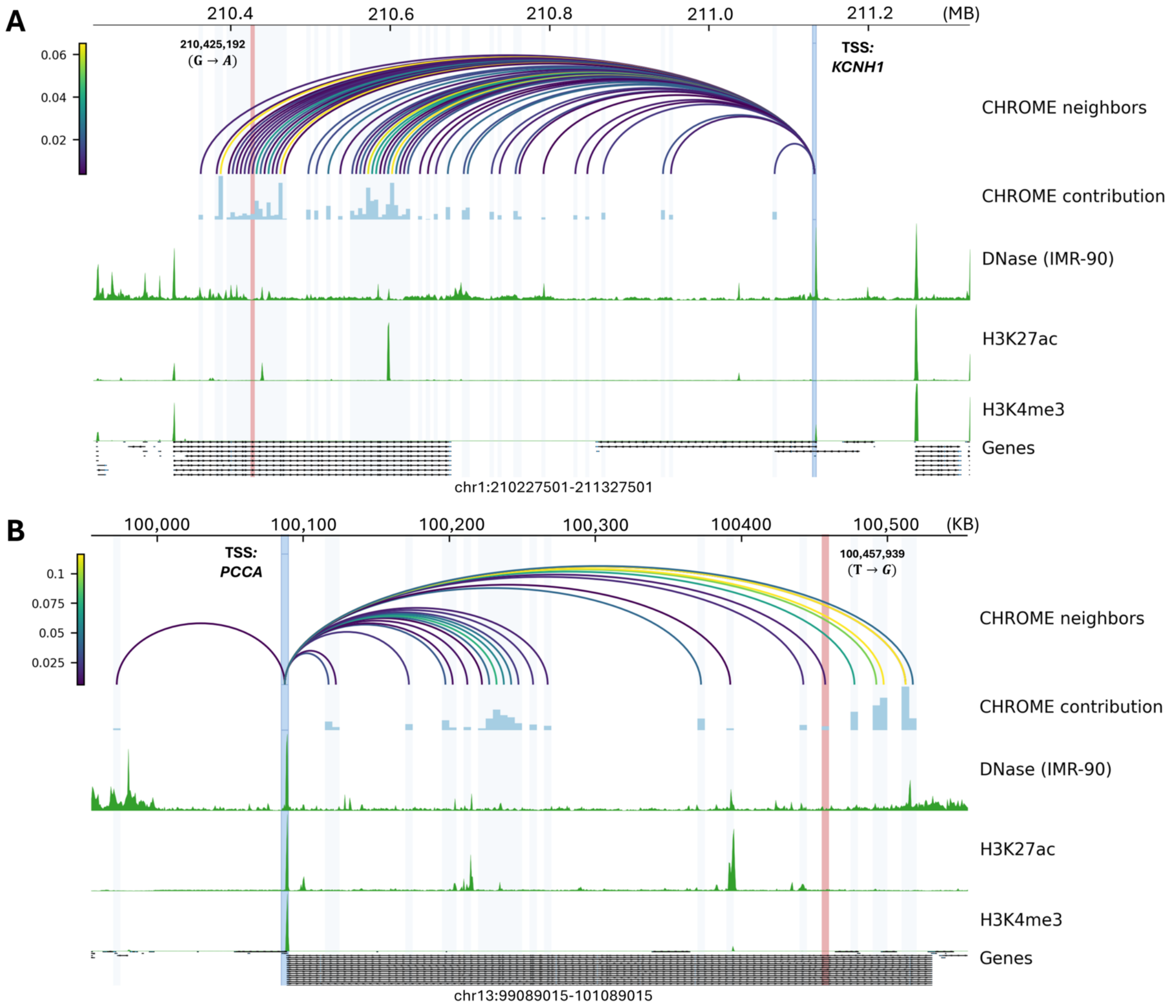


**Supplementary Figure S10: Case studies of eQTL-containing neighbor contributions to TSS-centered loci in Lung (IMR-90).** CHROME neighbor–contribution maps are shown for two representative loci in IMR-90 (mapped to GTEx Lung), where the *eQTL resides in a spatially connected non-random Hi-C neighbor loci* that contributes to the TSS-centered 5 kb locus. **(A)** *KCNH1*: a variant (G→A) at chr1:210,425,192 lies in a distal non-random locus; it contributes to the TSS-centered 5 kb locus, with the contributing neighborhood extending up to ∼0.8 Mb from the center. **(B)** *PCCA*: a variant (T→G) at chr13:100,457,939 lies in a non-random locus; contributions to the TSS-centered 5 kb locus span a more compact region of ∼0.5 Mb. Tracks shown include CHROME neighbor arcs, CHROME contribution scores, DNase accessibility, histone modifications (H3K27ac, H3K4me3), and annotated gene models.

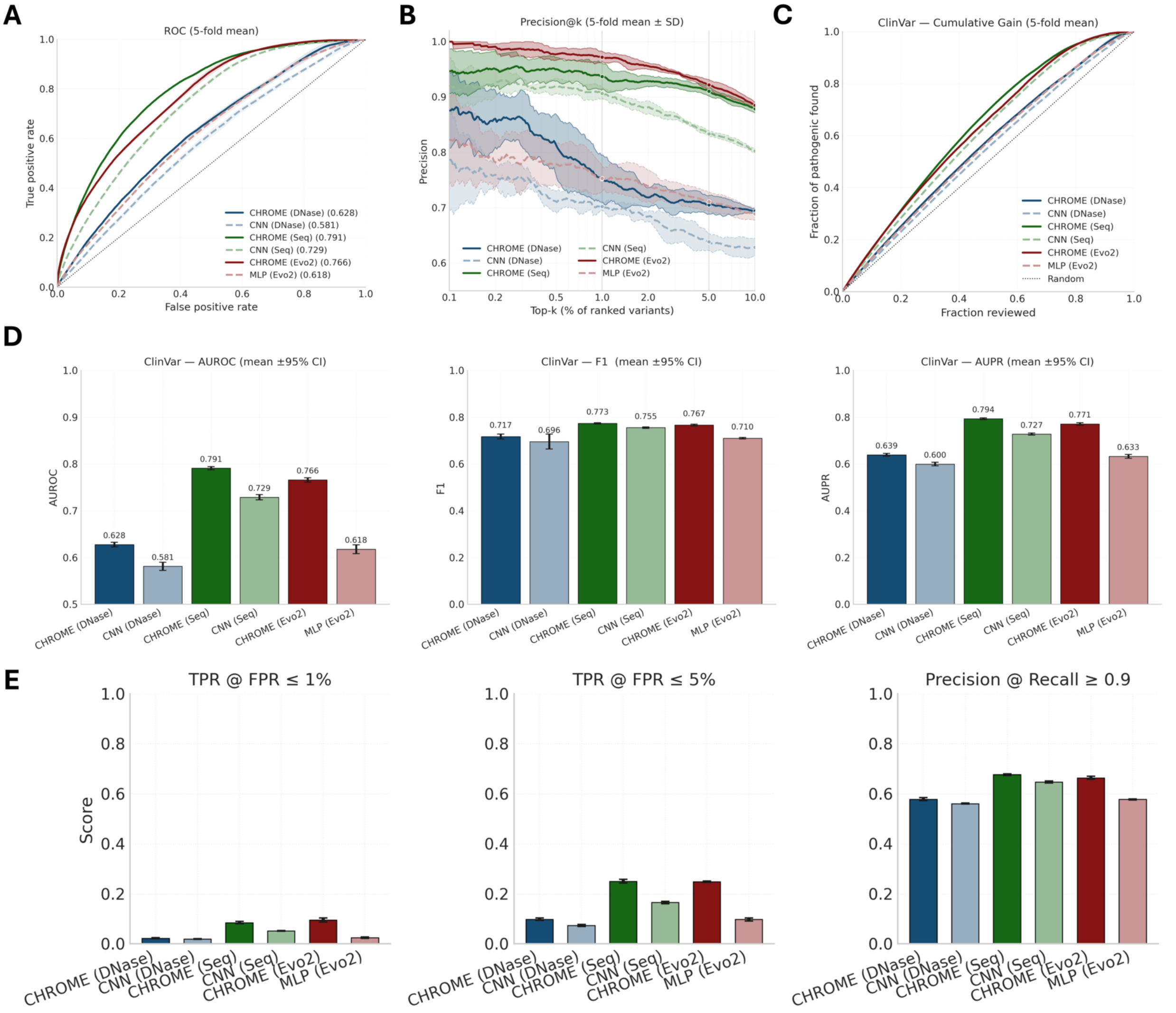


**Supplementary Figure S11: Supplementary evaluation of ClinVar pathogenicity prediction.** **(A)** ROC curves (5-fold mean); CHROME models (solid lines) generally achieve higher AUROC than CNN/MLP baselines (dashed), with pronounced gains for sequence- and Evo2-based inputs. **(B)** Precision@k (5-fold mean ± SD), showing that graph-based CHROME maintains higher precision among top-ranked variants across a broad range of $k$. **(C)** Cumulative gain curves indicating stronger enrichment of pathogenic variants among top-ranked predictions relative to baselines and random expectation. **(D)** Summary bar plots (mean ± 95% CI) for AUROC, F1, and AUPR across folds. **(E)** Operating-point analyses: (left) TPR at FPR $\leq 1\%$, (middle) TPR at FPR $\leq 5\%$, and (right) precision at recall $\geq 0.9$, where CHROME exhibits stronger or comparable performance at clinically relevant thresholds.

**Supplementary Table S1: Per-chromosome concordance between ABC and CHROME contributions.** For GM12878 and K562, Spearman correlations are computed between ABC Activity-by-Contact scores and CHROME (DNase) neighbor-to-center contributions on a per-chromosome basis. We report Spearman_all using all evaluated links and Spearman_thr after restricting to links with ABC score > 0.015.

| Chromosome | GM12878 | | K562 | |
|---|---|---|---|---|
| | Spearman_all | Spearman_thr | Spearman_all | Spearman_thr |
| chr1 | 0.048 | 0.023 | 0.193 | 0.020 |
| chr2 | -0.023 | -0.025 | 0.206 | 0.052 |
| chr3 | -0.038 | -0.053 | 0.199 | 0.068 |
| chr4 | -0.008 | 0.064 | 0.227 | 0.088 |
| chr5 | -0.040 | 0.014 | 0.275 | 0.046 |
| chr6 | 0.019 | -0.009 | 0.156 | 0.066 |
| chr7 | 0.050 | 0.004 | 0.156 | -0.078 |
| chr8 | -0.007 | 0.077 | 0.227 | 0.078 |
| chr9 | -0.015 | 0.150 | 0.232 | 0.056 |
| chr10 | -0.053 | -0.008 | 0.228 | 0.026 |
| chr11 | 0.011 | 0.001 | 0.205 | 0.011 |
| chr12 | 0.018 | 0.036 | 0.242 | 0.052 |
| chr13 | 0.009 | 0.141 | 0.271 | 0.001 |
| chr14 | -0.022 | 0.002 | 0.217 | -0.061 |
| chr15 | -0.024 | -0.065 | 0.195 | -0.008 |
| chr16 | 0.027 | 0.021 | 0.195 | -0.030 |
| chr17 | -0.064 | -0.007 | 0.192 | -0.025 |
| chr18 | -0.010 | -0.060 | 0.145 | 0.043 |
| chr19 | 0.017 | 0.001 | 0.183 | 0.011 |
| chr20 | -0.071 | -0.001 | 0.140 | -0.100 |
| chr21 | -0.018 | -0.218 | 0.219 | -0.010 |
| chr22 | -0.014 | 0.020 | 0.298 | 0.086 |
| chrX | 0.047 | 0.118 | 0.172 | -0.024 |

**Supplementary Table S2: Baseline CNN and MLP encoder architectures.** All models predict 751 ChIP-seq profiles with a final linear layer.

| Baseline architectures | |
|---|---|
| **CNN (seq)** | **Input:** one-hot DNA (4 channels), 5 kb window per locus |
| | 1D Convolution (kernel size: 10, in: 4, out: 256), Dropout: 0.1 |
| | 1D Convolution (kernel size: 10, out: 256), Max-pool (size: 5), BatchNorm, Dropout: 0.1 |
| | 1D Convolution (kernel size: 8, out: 360), Dropout: 0.1 |
| | 1D Convolution (kernel size: 8, out: 360), Max-pool (size: 4), BatchNorm, Dropout: 0.1 |
| | 1D Convolution (kernel size: 8, out: 512), Dropout: 0.2 |
| | 1D Convolution (kernel size: 8, out: 512), BatchNorm, Dropout: 0.2 |
| | Global average pooling |
| | Linear projection (to 256) $\rightarrow$ ReLU $\rightarrow$ BatchNorm |
| | Linear classifier (256 $\rightarrow$ 751) |
| **CNN (DNase)** | **Input:** DNA one-hot (4) + DNase (1) $\Rightarrow$ 5 input channels |
| | 1D Convolution (kernel size: 10, in: 5, out: 256), Dropout: 0.1 |
| | 1D Convolution (kernel size: 10, out: 256), Max-pool (size: 5), BatchNorm, Dropout: 0.1 |
| | 1D Convolution (kernel size: 8, out: 360), Dropout: 0.1 |
| | 1D Convolution (kernel size: 8, out: 360), Max-pool (size: 4), BatchNorm, Dropout: 0.1 |
| | 1D Convolution (kernel size: 8, out: 512), Dropout: 0.2 |
| | 1D Convolution (kernel size: 8, out: 512), BatchNorm, Dropout: 0.2 |
| | Global average pooling |
| | Linear projection (to 256) $\rightarrow$ ReLU $\rightarrow$ BatchNorm |
| | Linear classifier (256 $\rightarrow$ 751) |
| **MLP (Evo2)** | **Input:** Evo2 embedding (4096-d) at the center locus |
| | Linear (4096 $\rightarrow$ 128), ReLU, Dropout: 0.5 |
| | Linear (128 $\rightarrow$ 128), ReLU, Dropout: 0.5 |
| | BatchNorm(128) |
| | Linear (128 $\rightarrow$ 751) |
| | Sigmoid (multi-label) |

**Supplementary Table S3: CHROME (graph attention over non-random Hi-C neighborhoods).** A CNN/MLP encoder produces node embeddings; two GAT layers aggregate non-random neighbors; pooled neighbor context is concatenated with the center node before the final classifier.

| CHROME architectures | |
|---|---|
| **CHROME (seq)** | **Node encoder (shared CNN on each node)** |
| | Input: DNA one-hot (4), 5 kb window |
| | Conv1D (k=10, in:4, out:256) → Dropout 0.1 |
| | Conv1D (k=10, out:256) → MaxPool(5) → BatchNorm → Dropout 0.1 |
| | Conv1D (k=8, out:360) → Dropout 0.1 |
| | Conv1D (k=8, out:512) → BatchNorm → Dropout 0.2 |
| | Global average pooling → Linear (to *embed_dim*) → ReLU → BatchNorm |
| | **Graph encoder (non-random Hi-C neighbors)** |
| | GATConv (*embed_dim* → *embed_dim*, heads=4, concat), ELU |
| | GATConv (4×*embed_dim* → *embed_dim*, heads=1), Dropout 0.2 |
| | Global mean pooling over neighbors ⇒ *neighbor_ctx* (*embed_dim*) |
| | Concatenate [*center_emb* ; *neighbor_ctx*] (2×*embed_dim*) |
| | Linear classifier (2×*embed_dim* → 751) |
| **CHROME (DNase)** | **Node encoder (shared CNN on each node)** |
| | Input: DNA one-hot (4) + DNase (1) ⇒ 5 channels, 5 kb window |
| | Conv1D (k=10, in:5, out:256) → Dropout 0.1 |
| | Conv1D (k=10, out:256) → MaxPool(5) → BatchNorm → Dropout 0.1 |
| | Conv1D (k=8, out:360) → Dropout 0.1 |
| | Conv1D (k=8, out:512) → BatchNorm → Dropout 0.2 |
| | Global average pooling → Linear (to *embed_dim*) → ReLU → BatchNorm |
| | **Graph encoder (non-random Hi-C neighbors)** |
| | GATConv (*embed_dim* → *embed_dim*, heads=4, concat), ELU |
| | GATConv (4×*embed_dim* → *embed_dim*, heads=1), Dropout 0.2 |
| | Global mean pooling over neighbors ⇒ *neighbor_ctx* (*embed_dim*) |
| | Concatenate [*center_emb* ; *neighbor_ctx*] (2×*embed_dim*) |
| | Linear classifier (2×*embed_dim* → 751) |
| **CHROME (Evo2)** | **Node encoder (shared MLP on each node)** |
| | Input: Evo2 embedding (4096-d) |
| | Linear (4096 → 128), ReLU, Dropout 0.5 |
| | Linear (128 → 128), ReLU, Dropout 0.5, BatchNorm(128) |
| | **Graph encoder (non-random Hi-C neighbors)** |
| | GATConv (128 → 128, heads=4, concat), ELU, Dropout 0.5 |
| | GATConv (4×128 → 128, heads=1), Dropout 0.5 |
| | Global mean pooling over neighbors ⇒ *neighbor_ctx* (128) |
| | Concatenate [*center_emb* ; *neighbor_ctx*] (256) |
| | Linear (256 → 751) |
| | Sigmoid (multi-label) |

### S1 Hi-C preprocessing and identification of non-random contacts

Hi-C contact matrices for GM12878, K562, IMR-90, and HepG2 were retrieved from the 4DN Data Portal [2, 3] and mapped to the hg38 reference genome. To identify physically specific non-random chromatin contacts, we constructed a self-avoiding polymer ensemble null model following approaches described in the CHROMATIX framework [4, 5]. Chromatin was modeled as a self-avoiding polymer chain, with each bead representing a 5 kb genomic locus. Ensembles of random polymers spanning 4 Mb (800 beads) were generated by Monte Carlo sampling within a confined nuclear volume [6, 7], independent of Hi-C measurements, thereby providing an unbiased background expectation. The 4 Mb window size was selected as a practical upper bound that encompasses typical TAD-scale structures while remaining computationally tractable. For each bead pair, a geometric contact was defined when the three-dimensional distance between bead centers was less than 80 nm [8]. Across $5 \times 10^5$ random polymer conformations, we tallied the frequency of each bead pair in contact to estimate the random contact probability at a given genomic separation, and applied bootstrap resampling on the null ensemble to construct distance-stratified distributions.

Measured Hi-C contact frequencies were then compared against this null distribution. For each genomic bin pair, a $p$-value was computed as the fraction of bootstrap-null frequencies greater than or equal to the observed Hi-C frequency. False discovery rate (FDR) was controlled at 5% using the Benjamini–Hochberg procedure [9], and pairs passing this threshold were designated as non-random contacts. These interactions represent chromatin contacts that occur more frequently than expected from random polymer collisions and thus reflect biologically specific spatial organization. The resulting set of non-random contacts was used to build CHROME graphs, ensuring that only physically specific interactions contribute to neighborhood construction in downstream modeling.

### S2 Chromatin accessibility data processing

Following the EPCOT framework [10], we processed DNase-seq datasets to generate cell line–specific chromatin accessibility profiles for CHROME. Raw DNase-seq BAM files were downloaded from the ENCODE [1] Project, and replicates for each cell line were merged prior to normalization. Merged BAMs were converted to bigWig format using deepTools [11] bamCoverage with RPGC normalization (binSize = 1), ensuring cross-sample comparability. This standardized workflow provides consistent accessibility signals across all cell lines used in CHROME.

DNase-seq accessibility datasets were obtained from ENCODE under the following accession numbers: GM12878 (ENCSR000EMT), K562 (ENCSR000EOT), IMR-90 (ENCSR477RTP), and HepG2 (ENCSR149XIL). The processed and normalized bigWig files were subsequently used as input tracks for cell line–specific feature extraction.

### S3 Cell line–specific ChIP-seq prediction

For the ChIP-seq prediction experiments, 1 kb genomic bins were labeled and expanded by 2 kb upstream and downstream, following a strategy similar to EPCOT [10], yielding 5 kb sequence windows that defined the center nodes. CHROMATIX [5] validated non-random Hi-C contacts connected to each center node were incorporated as leaf nodes [12], forming the local CHROME signal graph.

We evaluated three types of input encodings: (1) one-hot representations of raw DNA sequence, (2) one-hot sequence augmented with cell line–specific DNase-seq accessibility signals, and (3) dense sequence embeddings derived from the Evo2 foundation model. For consistency, only the forward DNA strand and DNase signal were used in all models.

Model training followed a chromosome-wise partition: chromosome 9 was held out for validation, chromosome 8 for testing, and the remaining autosomes (excluding chromosome Y) were used for training. Only bins containing at least one positive epigenomic signal in the center were included in any split. Consistent with EPCOT [10], which reported similar predictive performance for ATAC-seq and DNase-seq accessibility, we used DNase-seq as the chromatin accessibility input to capture cell line–specific context.

#### S4 Model encoders and graph attention framework

As summarized in Supplementary Tables S2–S3, all baseline and CHROME variants share a consistent encoder–decoder organization. The CNN encoders process 5 kb loci using one-dimensional convolutional filters (kernel size = 19, stride = 1) [13, 14] followed by max-pooling and two fully connected layers. The DNase variant concatenates normalized accessibility values as an additional input channel, while the Evo2 variant replaces the convolutional front end with a 4096-dimensional embedding projected through a lightweight MLP. Graph attention layers (two hops, 8 attention heads, hidden dimension = 128, dropout = 0.1) propagate encoded features across non-random chromatin contacts, enabling spatial information flow up to 4 Mb. Global mean pooling aggregates subgraph representations for multi-label prediction of 751 ChIP-seq assays per cell line. These components collectively form a modular architecture in which encoder type determines feature modality and the GAT trunk captures 3D chromatin context.

### S5 Contribution extraction and interpretation

All contribution analyses were based on the CHROME (DNase) model, which showed the highest predictive performance across cell lines. For each genomic graph, neighbor-to-center contributions were derived from the attention coefficients of the final GAT layer[12, 15]. For an edge connecting a neighbor node $i$ to the center node $j$, the raw contribution was defined as

$$c_{ij}^{\text{raw}} = \alpha_{ij} \cdot \|h_i\|_1,$$

where $\alpha_{ij}$ denotes the learned attention coefficient and $h_i$ is the hidden feature representation of node $i$ after the first GAT layer. These raw values capture the absolute influence of each neighbor on the center, integrating both its attention weight and activation strength.

To enable within-locus comparison, we also computed normalized contributions:

$$\tilde{c}_{ij} = \frac{c_{ij}^{\text{raw}}}{\sum_{k \in \mathcal{N}(j)} c_{kj}^{\text{raw}}},$$

such that all incoming contributions to a given center node sum to one. The normalized $\tilde{c}_{ij}$ therefore represents the fractional regulatory importance of neighbor $i$ relative to all neighbors of the same locus, while $c_{ij}^{\text{raw}}$ retains the absolute influence magnitude.

In subsequent analyses, the two forms were used for complementary purposes. Raw contributions were used to evaluate distance-dependent decay and to summarize genome-wide distributions of neighbor effects as a function of genomic separation. Normalized contributions were used for comparisons against Activity-by-Contact (ABC) scores [16] and for locus-level visualizations (e.g., NDUFB6, RGS3, KCNH1, and PCCA), as they reflect relative contribution shares within each local chromatin neighborhood. Values were averaged across attention heads to obtain a single per-edge contribution estimate.

### S6 eQTL analyses

We applied CHROME to predict tissue-specific expression quantitative trait loci (eQTLs) using GTEx [17] v8 fine-mapped variants curated in Borzoi [18]. Variants from lymphoblastoid and lung tissues were mapped to the corresponding GM12878 and IMR-90 cell lines, respectively. Each eQTL-containing 5 kb locus was used as the center node, and all non-random Hi-C contacts connected to this locus were included as neighboring nodes to construct a variant-centered graph. In total, 314 variant-centered graphs were constructed for GM12878 and 1,299 for IMR-90, each supported by at least one non-random contact within 4 Mb.

For each variant, CHROME generated a graph embedding from the non-random neighborhood using models trained on ChIP-seq prediction (transfer setting), while a local baseline encoder produced a dense embedding from the target gene's TSS-centered 5 kb locus. These two embeddings were concatenated and passed to a multilayer perceptron (MLP) to classify whether the variant–gene pair corresponds to a true eQTL association.

To further assess whether CHROME captures the directionality of eQTL effects, we used the CHROME (DNase) model to estimate the variant-induced change in regulatory profiles. For each variant, both reference and alternate alleles were propagated through the CHROME (DNase) model, and the predicted differences ($\Delta$probabilities) across expression-associated ChIP-seq profiles were computed. These predicted shifts were then compared with the GTEx eQTL slope direction (positive or negative) to evaluate concordance between predicted regulatory perturbation and observed transcriptional change.

### S7 ClinVar analyses

ClinVar data [19] were obtained from the GRCh38 release of July 2024. A total of 212,542 entries, including both single, nucleotide variants and short indels annotated as benign or pathogenic, were retained after restricting to loci whose 5 kb loci contained at least one non-random Hi-C contact. To construct a consensus regulatory map, significant Hi-C contacts were intersected across GM12878, IMR-90, and K562, while DNase accessibility profiles were averaged over the same cell lines to approximate a cell-type–agnostic chromatin landscape. Variant-centered graphs were then assembled from these consensus maps.

Pathogenic enrichment analyses were performed using the ncVarDB [20] benchmark, with mutation-induced probability changes ($|\Delta \text{prob}|$) computed between mutant and reference alleles using the CHROME (DNase) model. Enrichment was evaluated separately for *cis* variants (located in the center locus) and *distal* variants (located in non-random loci). Both categories exhibited strong enrichment for pathogenic variants within the top 1% of $|\Delta \text{prob}|$ values, while distal variants consistently showed higher aggregated effects, suggesting that long-range chromatin contacts amplify the functional impact of pathogenic non-coding mutations.
